# RecA and RecB: probing complexes of DNA repair proteins with mitomycin C in live *Escherichia coli* with single-molecule sensitivity

**DOI:** 10.1101/2021.10.04.463068

**Authors:** Alex L. Payne-Dwyer, Aisha H. Syeda, Jack W. Shepherd, Lewis Frame, Mark. C. Leake

## Abstract

The RecA protein and RecBCD complex are key bacterial components for the maintenance and repair of DNA. RecBCD is a helicase-nuclease that uses homologous recombination to resolve double-stranded DNA breaks. It also facilitates coating of single-stranded DNA with RecA to form RecA filaments, a vital step in the double-stranded break DNA repair pathway. However, questions remain about the mechanistic roles of RecA and RecBCD in live cells. Here, we use millisecond super-resolved fluorescence microscopy to pinpoint the spatial localization of fluorescent reporters of RecA or RecB at physiological levels of expression in individual live *Escherichia coli* cells. By introducing the DNA crosslinker mitomycin C, we induce DNA damage and quantify the resulting steady state changes in stoichiometry, cellular protein copy number and molecular mobilities of RecA and RecB. We find that both proteins accumulate in molecular hotspots to effect repair, resulting in RecA stoichiometries equivalent to several hundred molecules that assemble largely in dimeric subunits before DNA damage, but form periodic subunits of approximately 3-4 molecules within mature filaments of several thousand molecules. Unexpectedly, we find that the physiologically predominant forms of RecB are not only rapidly diffusing monomers, but slowly diffusing dimers.

## 1. Introduction

Accurate duplication of the genome is crucial in all organisms, accomplished by a sophisticated molecular machine known as the replisome [1]. An inability to accurately replicate genetic material can lead to cell death and/or cancers [2,3]. Mitomycin C (MMC) is a naturally occurring antibiotic that can be used to controllably disrupt DNA replication, and thus a valuable reagent in studying DNA repair processes. It is used as a chemotherapeutic in treating several cancers [4] and retinopathies [5], and acts by targeting DNA deoxyguanosine (dG) residues [6], forming intrastrand or interstrand crosslinks [7]. If unrepaired, these structures can interfere with cellular processes such as transcription and replication, potentially leading to genome instability [8]. An encounter between a mitomycin C-induced crosslink and an approaching replisome may result in replisome disassembly and eventually a double strand break (DSB) [9]. RecBCD recognises DSBs in *E. coli* [10], processing the ends to generate 3’-ended single-stranded DNA (ssDNA) as a landing pad for the principal recombination protein, RecA [10]. Recombination of RecA-ssDNA complexes with the homologous DNA restores the replication fork, on which the replisome can be reloaded. The replisome may resume replication if the blocking adduct is repaired [11]. As a complex of individual RecB, RecC and RecD proteins, RecBCD is a versatile helicase-nuclease and underpins two major pathways for homologous DNA recombination, essential for DSB repair [10]. RecBCD activities involve several processes - it recognises and binds DSBs, begins unwinding both DNA strands, and also degrades both [10]. This activity continues unhindered until it encounters an octameric Chi site that induces a shift in enzyme activity to degrade only the 5’-ended strand [12,13]. This activity shift results in a 3’-ended ssDNA overhang that facilitates RecA loading. A key function of RecA is its ability to form nucleoprotein filaments on exposed ssDNA in response to damage [14]. These filaments can infiltrate an intact duplex and, on finding homology, recombine with the infiltrated duplex [15,16]. The extension of filaments along the cell accelerates this homology search in a non-linear fashion [17]. Following further processing of the resulting structure, primosome proteins establish an intact replisome thereby enabling replication to resume [18]. Recombination proteins, such as RecBCD, need access to replication-transcription conflict sites and collapsed forks, but if RecBCD is missing then double-stranded DNA (dsDNA) is degraded by exonucleases [19,20], possibly resulting from replisome disassembly. However, how RecA stabilizes blocked forks remains an open question.

The nucleoprotein filaments formed by RecA are both a requisite and a hallmark of the cell-wide SOS response [21–24]. The SOS response is a regulatory shift that promotes cell survival in adverse conditions associated with increased rates of interrupted replication and DNA damage [25]. The SOS response to DNA damage induced by antimicrobials plays a major role in the emergence of persister cells [26] and wider antimicrobial tolerance on a population level [27].

Given these far-reaching implications of RecA and RecB activity as studied comprehensively with mutants [12,24,28–32], it is important to establish the number of molecules present in cells, how they are spatially distributed and organized, and how these are affected by antimicrobials such as MMC. Here, we use millisecond super-resolved Slimfield microscopy [33] in live *E. coli* containing genomically-encoded fluorescent fusions RecA-mGFP [34] and RecB-sfGFP [35]. Since RecA fusion constructs retain only partial function, our approach makes use of a merodiploid RecA fusion that expresses from one copy of the native gene and one copy of the *recA4155* fusion construct [34]. This strain rescues approximately wild-type sensitivity with mixed assemblies of the two RecA proteins [34].

We use Slimfield microscopy to visualise the spatial distribution of RecA and RecB fluorescent proteins in individual cells. From these quantitative images, we identify diffraction-limited local intensity maxima (we denote these as *foci –* see Table 1 for a description of technical Slimfield microscopy nomenclature used in this study) to a lateral spatial precision of 40 nm [36]. Slimfield uses ~millisecond sampling that is sufficiently rapid to link the moving foci derived from the same emitter sources over sequential image frames, following appropriate bespoke particle tracking analysis [33,37,38], into *tracks*. Each of these tracks implies the presence of a particle containing one or more associated molecules; typically more than one prior to photobleaching, so more generally, we term each a *molecular assembly*. These tracks reveal the detailed diffusion of labelled RecA and RecB assemblies in the cytoplasm of a living cell. By using the single-molecule sensitivity of Slimfield microscopy, we are able to quantify single-molecule photobleaching steps in fluorescence intensity, to identify the characteristic brightness of a single fluorescent protein [33]. Not only does this calibration apply to the fraction of the fluorescence intensity for each tracked assembly, but also to the GFP fluorescence in the whole, or part, of each cell. We use this to determine the number of GFP-labelled molecules within each tracked assembly (the *stoichiometry*), and the total number of fluorescently-labelled molecules within each cell (the *cellular protein copy number*), or intracellular segment (the *segment protein copy number*). Those fluorescent molecules which contribute to the copy number above the cell’s autofluorescent background but are not detected as foci (typically due to high, uniform emitter density and/or excessive mobility) are denoted the *pool*.

**Table 1.** Definitions of quantitative analysis metrics for Slimfield.

| **Metric / Object** | **Definition** |
| --- | --- |
| *Segment* | An area of the image defined by a contiguous subset of pixels in a binary mask. This area either corresponds to a whole cell (a *cell mask*), or more typically a region inside the cell (an *intracellular segment*) of high fluorescent intensity. The term “*segment*” refers to an intracellular segment unless otherwise stated. |
| *Cell mask* | A segment containing the outline of one cell. These are extracted using a machine learning protocol (Supplementary Methods). |
| *Intracellular segment* | A segment inside the cell. These are extracted from the set of foci localized in that cell by rendering a superresolved image, followed by local Otsu thresholding (Materials and Methods 4.3.5), with the intention of isolating RecA objects that resemble nucleoprotein filaments or bundles. |
| *Assembly* | A group of labeled molecules physically associated with one another, either directly or indirectly, such that their diffusive movement is strongly correlated, and therefore always detected in the same track. |
| *Focus (foci)* | A spot-like local intensity maximum in a single frame, which corresponds to a localized group of labeled molecules (Materials and Methods 4.3.1). Associated properties include centroid location, total intensity, and signal-to-noise ratio. |
| *Track* | A set of foci in adjacent frames that are spatially close enough to form a contiguous trajectory (Materials and Methods 4.3.1). Typically associated with a single molecular assembly, or a group of strongly colocalized assemblies. |
| *Diffusion coefficient* | Measure of the random microscopic motion of a specific track based on the increase in the mean square displacement of its intensity centroid over time (Materials and Methods 4.3.2). |
| *Characteristic single-molecule brightness* | The average sum of pixel values in foci associated with a single fluorescent reporter molecule (e.g. mGFP), under a fixed imaging condition (Materials and Methods 4.3.3). Equivalent to the modal step size in intensity for tracks in the final stage of photobleaching (Figure S1). |
| *Stoichiometry* | The number of fluorescently labeled molecules in a specific track. This is estimated by extracting the sequence of foci belonging to that track, then extrapolating the sum of pixel values in each focus backwards along that sequence to get an initial track intensity that is independent of photobleaching (Materials and Methods 4.3.4). The initial track intensity is then divided by the characteristic single-molecule brightness. |
| *Periodicity* | The population-averaged number of fluorescently labeled molecules in inferred repeat units within tracked objects. Estimated by averaging the consistent intervals between nearest-neighbor peaks in the population-level stoichiometry distribution (Materials and Methods 4.3.5). |
| *Integrated intensity* | The total fluorescence intensity of a segment in pixel counts, normalised by the characteristic single molecule brightness (Materials and Methods 4.3.6-7). |
| *Cellular (or segment) protein copy number* | The average number of molecules in a cell (or intracellular segment), as estimated from the increase in integrated intensity above negative control (i.e. subtracting the contribution from autofluorescence (Materials and Methods 4.3.6-7). |
| *Pool* | The intracellular fluorescence which is not detected in tracks. |
| *Pool stoichiometry* | The number of untracked, labeled molecules within an area of the pool equal to the size of one diffraction-limited focus (Materials and Methods 4.3.6). |

**Table 2.**
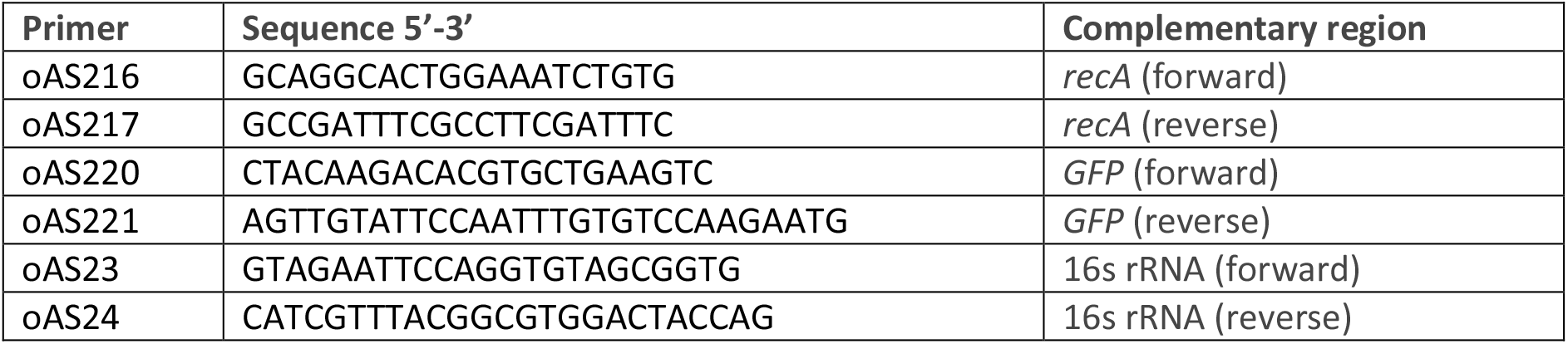
Primers used for qPCR to quantify mRNA of *recA, recA-gfp* and housekeeping gene *rrsA*.

Slimfield has some similarities to single particle tracking photoactivation localization microscopy (sptPALM) [39–41], however, our approach is simpler, requiring only constitutively expressed fluorescent reporters such as GFP, and trades off the condition of observing exclusively single molecules in order to measure the stoichiometry of dynamic assemblies far more accurately. This is a deliberate advantage of our technique over other single-molecule microscopy techniques as previously used to count RecB content in cells molecule-by-molecule [35].

Prior to MMC treatment, only point-like assemblies of RecA or RecB are detectable. RecA presents far brighter fluorescence in a cell than RecB, indicating both a typical stoichiometry and a cellular protein copy number that are 2-3 orders of magnitude greater. On treatment with MMC, we observe an increase in the average cellular protein copy number of RecA, but not of RecB, in each cell, with up to 20% of cells devoid of RecB assemblies. MMC induces the formation of RecA assemblies larger than can be captured in single foci, and we interpret these as RecA nucleoprotein filaments, or bundles of filaments [24,30,42–47], typically associated with the SOS response.

Between cellular states of SOS readiness and MMC-induced response, the stoichiometries of RecA assemblies increase, and the diffusion coefficients of assemblies decrease correspondingly. We also discover surprisingly consistent intervals between the stoichiometries of different assemblies in each condition. We interpret the average number of molecules in the intervals (the *periodicity*) as indicative of an oligomeric structural repeat unit that comprises assemblies. The periodicity of RecA assemblies changes from dimeric in character to groups of roughly 3-4 molecules in response to MMC, while the periodicity of RecB assemblies is dimeric, and insensitive to MMC treatment.

Our results shed new light on the relations between structure and function for RecA and RecBCD in mediating repair upon DNA damage.

## 2. Results

### 2.1 Abundance of RecA, but not RecB, increases on MMC-induced DNA damage

We first optimised MMC treatment conditions so that they did not cause cellular filamentation in wild type cells (Materials and Methods 4.1, Figure S1) but did induce the SOS response [48], since cells would then be sensitised to MMC if the SOS response is blocked [48]. Filamentation and loss of viability was also minimal for the labeled strains (Figure 1), hence we used the same MMC treatment for all strains. Given that SOS induction in these and related strains typically takes <20 min [34,48], the kinetics of initial SOS induction due to MMC will likely reach steady state within the 180 min MMC exposure that we used. In light of the timescale of the initial SOS induction, RecA or RecB dynamics were not in the scope of our study here, but rather the steady state effect of MMC on the distribution and molecular organization of RecA and RecB.

**Figure 1.**
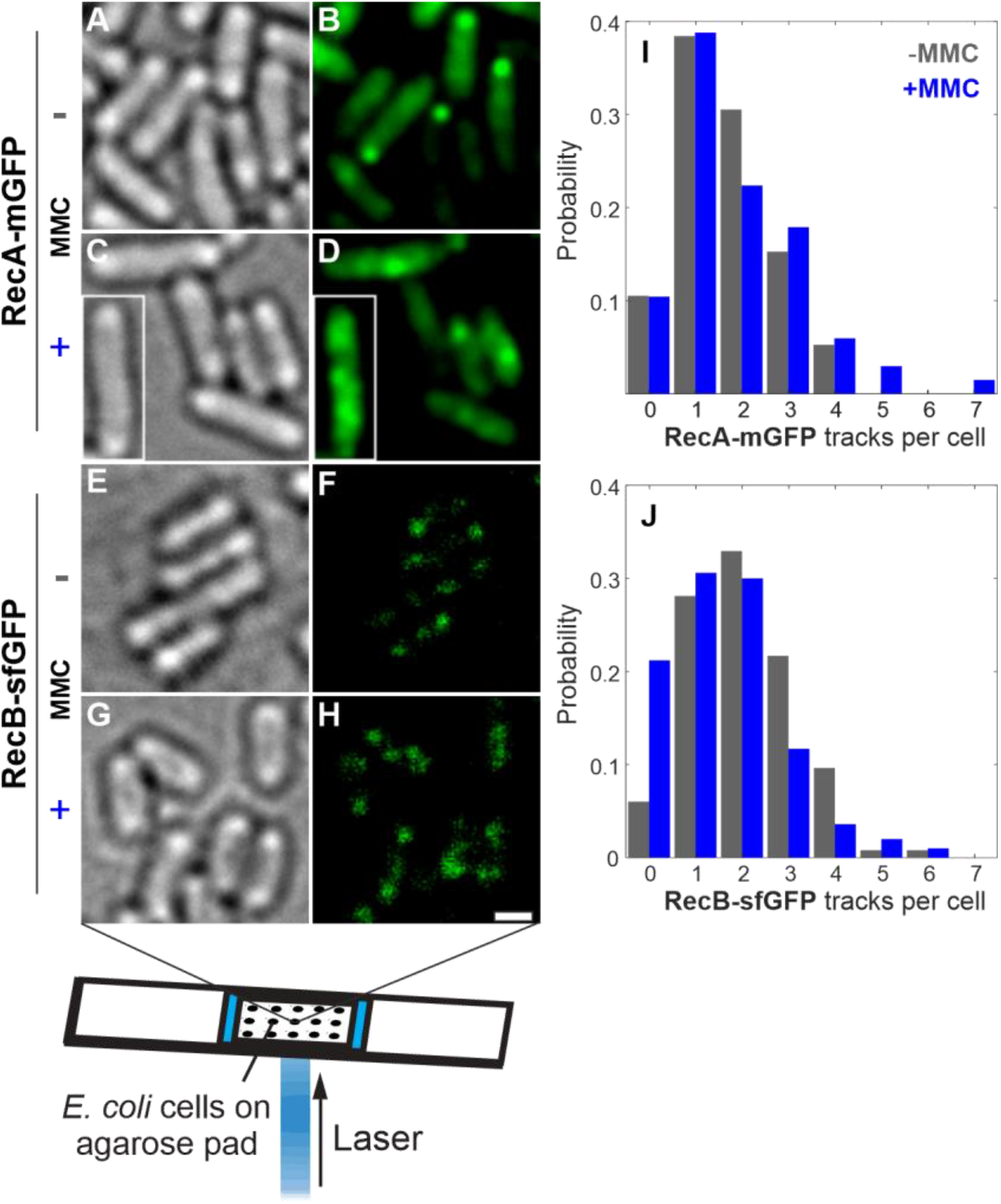
Brightfield and Slimfield (mean average of 3 initial frames) of live *E. coli* in 56-salts minimal media, labeled at RecA-mGFP or RecB-sfGFP before and after MMC treatment. Inset (C,D) is another cell transplanted from the same acquisition outside the cropped field of view at the same scale. Brightness of RecB-GFP Slimfield panels (F,H) scaled 100× *vs*. RecA-mGFP panels (B,D). Scale bar 1 μm. (I,J) Probability distributions for number of tracks detected per cell. Tracks are identified in post-acquisition analysis (Materials and Methods 4.3.1) by first detecting foci as local fluorescent maxima, then linking nearest-neighbour foci in subsequent frames.

We performed brightfield and Slimfield microscopy in each field of view (Materials and Methods 4.2). Binary masks for each cell were extracted independently of any fluorescence signal, using a machine learning segmentation protocol on brightfield images. We then applied these masks to fluorescence images to eliminate extracellular background and facilitate statistics on a cell-by-cell level (Figure S5 and Supplementary Methods). Using home-written automated particle tracking and analysis software ADEMScode [49], we identified fluorescent foci from local intensity maxima in each cell and linked these into tracks (Materials and Methods 4.3.1). We determined the stoichiometry (the number of molecules present) of each track, from the summed pixel intensity values corresponding to the start of each track before any photobleaching has occurred (Materials and Methods 4.3.4), normalized by brightness corresponding to a single molecule of GFP (Materials and Methods 4.3.3).

From the localizations of foci during photobleaching, we also reconstructed superresolved images of fluorescent RecA structures. We extracted binary masks from highly fluorescent regions of interest in these images (denoted *intracellular segments*, or simply *segments* for brevity) using a classical segmentation method (Materials and Methods 4.3.7), which enabled statistics on an intracellular segment level (Figure 3 and Figure S3).

Separately, from the cell masks (or intracellular segments) we also calculated the cellular (or segment) protein copy number (Materials and Methods 4.3.6-7); first we summed the pixel values in each cell or segment area and normalized these by the characteristic brightness of a single GFP to obtain the total intensity within that region, expressed in molecules [50]. Taking the difference from an equivalent area of the control strain that does not express GFP, then yields the cellular or segment protein copy number corrected for any cellular autofluorescence.

Since the RecA-mGFP strain is merodiploid, both the *recA-mgfp* gene fusion construct and the unlabeled endogenous *recA* gene are expressed simultaneously [34]. However, their expression levels are not necessarily identical, nor equivalently inducible by MMC. From previous estimations of the relative *lexA* suppression rates of the relevant *recA* promoters [51], reasonable expectations are that a majority of the RecA present in the cell will be labelled with mGFP, and that RecA-mGFP is 2-3 fold less inducible under the SOS response as endogenous RecA [52]. We estimated the different cellular levels of unlabeled RecA *vs*. RecA-mGFP using Western blotting (Figure S4), which confirmed that the RecA-mGFP was in excess compared to the endogenous protein before and after treatment. Both qPCR and Western blots indicated that both endogenous RecA and RecA-mGFP are inducible by MMC treatment (Materials and Methods 4.4), with the RecA-mGFP indeed about half as inducible (Figure S4). Therefore, the total (i.e. labelled plus unlabelled) amount of RecA protein present, whether as stoichiometry, periodicity or protein copy numbers, is higher than that reported for the RecA-mGFP data directly, by an approximate correction factor of 1.3-fold in the presence of MMC. In the absence of MMC the relative amount of RecA-mGFP to RecA is large enough that the correction factor is effectively 1. As these corrections are indicative, we do not apply them in the early stages of the Results, but present them later only where relevant to interpretations (Results 2.3 and Discussion).

We find that in the absence of MMC, RecA-mGFP has an an approximately uniform distribution in the cytoplasm that is occasionally punctuated by bright fluorescent foci that can be linked into tracks (Figure 1B). The cellular protein copy number of RecA-mGFP increases from 11,400 ± 200 molecules (±SEM) in untreated cells to 19,500 ± 300 molecules in MMC treated cells (Figure S5A). MMC treatment resulted in the subset of these RecA-mGFP molecules that are localized in tracks (*i.e*., the mean summed stoichiometry of all tracks detected in the whole cell) approximately doubling from 510 ± 30 to 1,080 ± 60 molecules per cell (Table S1). We denote the fluorescently detected, but untracked, molecules of RecA as residing in a *pool*. The pool typically comprises molecules that are sufficiently dim, out-of-focus, and/or rapidly diffusing to evade direct particle-tracking-based detection; here the RecA concentration is exceptionally high such that the stochastic fluctuations corresponding to motion of discrete foci are partly averaged out. During photobleaching, the density of foci decreases, overlap decreases and tracks become more evident. The proportion of RecA-mGFP molecules in tracks is relatively low compared to the pool, but remains representative of the population of assemblies containing RecA-mGFP.

RecB-sfGFP also exhibited fluorescent tracks against a relatively diffuse background, before and after MMC treatment (Figure 1F,H). RecB-sfGFP foci were observed more commonly near the poles of the cell regardless of MMC (Figure 1G-H). Since the RecB-sfGFP fluorescence signal is comparatively small, estimates based on cellular protein copy number must account carefully for autofluorescence due to native components other than GFP. We estimate that the contribution of autofluorescence from the summed pixel intensity values from unlabeled MG1655 parental cells grown and imaged under identical conditions. We find that the mean level of RecB-sfGFP fluorescence was almost three times greater than the cellular autofluorescence (Figure S5B), therefore there is a comparatively large population of the cellular RecB-sfGFP that evades direct particle-tracking-based detection (*c.f*., slower sampled images from commercial confocal/epifluorescence microscope systems) and thereby comprise a RecB pool.

The cellular protein copy number of RecB-sfGFP does not decrease significantly following MMC-induced DNA damage, comprising 126 ± 11 molecules per cell before treatment and 101 ± 14 molecules following MMC treatment (Figure S5B, Brunner-Munzel (BM) test, n=246, p=0.0216 | NS, not significant at Bonferroni-adjusted α = 0.01). However, the mean number of RecB-sfGFP localized into tracks does decrease with MMC; just 13.6 ± 0.5 molecules per cell in all tracks, decreasing to 9.3 ± 0.3 on MMC treatment (BM test, n=246, p<0.001). This is clearly much smaller absolute number of tracked molecules per cell compared to RecA-mGFP, but a similar proportion of the cellular protein copy numbers (ranging from 6-10% in each case). The complementary fractions of the total RecA-mGFP and RecB-sfGFP molecules assigned to their respective pools are thus consistently high (89-95%). In the respective strains, the total concentration of RecB-sfGFP is much lower than that of RecA-mGFP, and this likely indicates the correspondingly more rapid diffusion of RecB-sfGFP species within the pool.

### 2.2 RecB forms characteristic puncta which are partially lost on MMC exposure

We detected typically 1-3 tracks of RecA-mGFP or RecB-sfGFP in each cell above the local background fluorescence (Figure 1I,J). However, each showed strongly opposing trends in the number of tracks observed upon MMC treatment. While RecA-mGFP showed no significant change in the mean number of tracks on MMC treatment (from 1.66 ± 0.06 to 1.86 ± 0.16 tracks per cell, BM test, n=60, p=0.50 |NS), MMC reduced the population average number of RecB-sfGFP tracks significantly, from 2.06 ± 0.09 to 1.56 ± 0.06 per cell.

If, however, we set aside the fraction of cells with no detected RecB-sfGFP tracks, the change in the mean number of RecB-sfGFP tracks is marginal, from 2.20 ± 0.07 to 1.98 ± 0.07 tracks (BM test, n=234, p=0.006). We see that the cells which continue to harbor RecB tracks are relatively unchanged by MMC, each containing an average of 12.1 ± 0.3 molecules per cell. The unexpected subset of cells that are devoid of RecB-sfGFP tracks increases from 6% to 21% of the population on MMC treatment. These otherwise resemble the other treated cells; rather than filamenting, they retain 92 ± 3% of the population averaged cell length and retain the same pool level of untracked RecB-sfGFP molecules.

The increase in the fraction of cells lacking RecB-sfGFP foci agrees with a model of random, independent survival of assemblies (Figure 1J, the MMC+ condition is consistent with Poisson distribution with same mean; Pearson χ^2^ test, dof=6, n=234, p=0.004).

RecA-mGFP foci were approximately two orders of magnitude brighter than those of RecB-sfGFP, corresponding to a greater apparent stoichiometry. A subset of polar assemblies in untreated cells are especially bright (Figure 2B); we defined this subset quantitatively by thresholding at 2× the mean stoichiometry of all assemblies. The mean stoichiometry of these bright assemblies is itself as high as 760 ± 40 molecules (Figure S3A).

**Figure 2.**
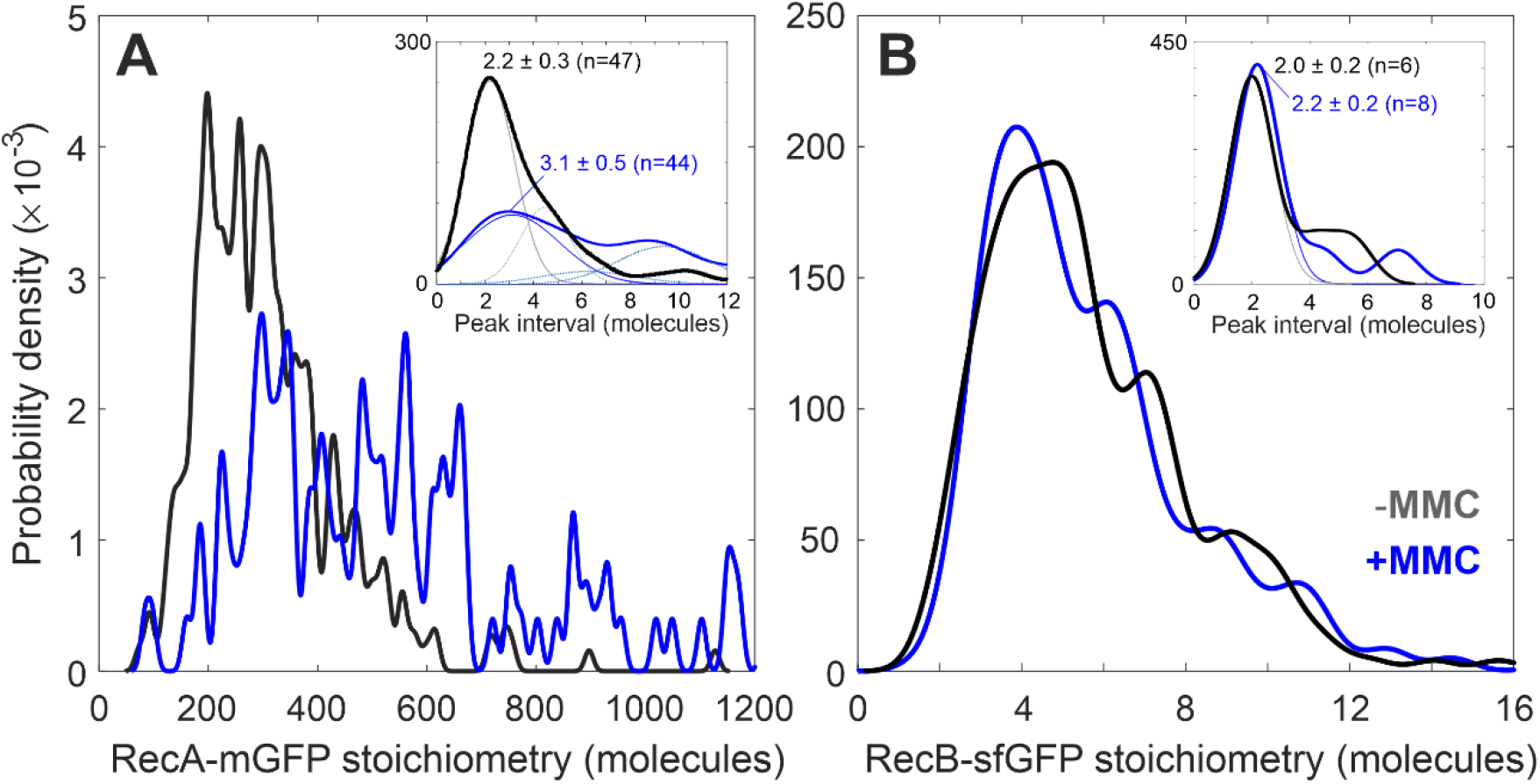
Stoichiometry distributions of detected foci of A) RecA-mGFP and B) RecB-sfGFP with (blue) or without MMC treatment (black), shown as kernel density estimates [53]. The statistics used for MMC-(MMC+) conditions include N=190 (67) whole RecA-mGFP cells containing n=316 (125) tracks, or whole N=249 (307) whole RecB cells containing n=514 (478) tracks within the cell masks. The use of ‘probability density’ reflects the fact that each distribution is continuous with a total area equal to 1, such that areas under the curve correspond to the probability that the stoichiometry of a given assembly falls within a range. The kernel width (the width for smoothing the discrete stoichiometry of each track) is 0.7 molecules following the known detection sensitivity to single GFP (A, inset and B, both panels), or 8 molecules for clarity (main panel A). Insets are the distributions of intervals between nearest neighbor stoichiometry peaks (solid curves) whose modal position, or periodicity, indicates the number of GFP-labeled molecules in a repeating subunit within molecular assemblies (Table 1). Overlaid are heuristic Gaussian fits that minimize a reduced χ^2^ metric, with components of equal width and whose centers are fixed at integer multiples to account for the detected optical overlap of an integer number of subunit repeats of tracked foci. The resulting fits comprise three components for RecA-mGFP with MMC treatment (blue, Pearson’s *R*^*2*^ = 0.979, 5 degrees of freedom (dof)) and two components for RecA-mGFP without MMC (grey, Pearson’s *R*^*2*^ = 0.961, 4 dof). The mode of the peak interval is indicated ±95% confidence interval, alongside the number of contributing peak pairs in the original stoichiometry distribution.

On treating with MMC, the RecA-mGFP mean stoichiometry almost doubled from 310 ± 8 to 580 ± 30 molecules per focus, reflecting further local accumulation of RecA-mGFP protein (Figure 2A). We find that most RecA-mGFP molecules comprise an untracked, diffusive pool, in which there are ~30 RecA-mGFP molecules in an area corresponding to that of a typical diffraction-limited focus (which we denote as the *pool stoichiometry*, Table 1). The fact that the relative increase in pool stoichiometry with MMC treatment to ~50 RecA molecules (Figure S5C) is smaller than the fractional increase in the amount of RecA in tracks (Figure 2), indicates that the MMC-driven upregulation of RecA disproportionately affects tracked assemblies. As such, either i) new assemblies are formed which contain much more RecA than those before MMC treatment, or ii) those assemblies that already contain local concentrations of RecA accumulate more RecA. Under our treatment protocol, these changes do not deplete the reservoir of RecA in the cytoplasm. This observation of localized accumulation of RecA-mGFP is consistent with prior reports of long nucleoprotein filament formation on single stranded DNA [34]. The increased number of RecA tracks we observe upon MMC treatment may therefore indicate greater occurrence of processed ssDNA.

The RecB-sfGFP mean stoichiometry decreases very slightly from 6.6 ± 0.1 to 6.1 ± 0.2 molecules per focus (BM, n=478, p<10^−6^) (Figure 2B). A mean of approximately 6 RecB-sfGFP molecules in each case can be explained if the assembly contains 3 identical subunits whose periodicity is 2 molecules (Figure 2B inset). A pool stoichiometry of ~1 molecule of RecB-sfGFP (Figure S5B,D) suggests that the untracked RecB-sfGFP are likely to be monomers irrespective of MMC treatment (BM test, n=243, p=0.27 | NS). We find that the untracked pool of RecB-sfGFP comprises 90 ± 1% of the total RecB-sfGFP molecules in the cell.

### 2.3. RecA reorganises into filaments with 3-4-mer subunits in response to MMC

RecA-mGFP and RecB-sfGFP stoichiometry distributions show clear and reproducible peaks (Figure 2A and 2B). One explanation is that each detected fluorescent focus has a diffracted-limited width of ~250 nm that may potentially contain more than one ‘subunit’ of RecA-mGFP or RecB-sfGFP, bound sufficiently to co-track, such that the measured focus stoichiometry may appear as an integer multiple of that subunit, manifest as periodic peaks on the focus stoichiometry distribution. The expected difference between pairs of values on the stoichiometry distribution is thus either zero or an integer multiple of the periodicity within measurement error. The magnitude of the most likely non-zero pairwise difference value corresponds to the periodicity, with less likely values corresponding to harmonic peaks. Our approach uses a modal estimate of the nearest-neighbor peak intervals (Materials and Methods 4.3.5), and therefore produces a continuous, heuristic estimate for the periodicity. We then compare this periodicity metric to realistic models with integer numbers of molecules. RecA-mGFP tracks have a periodicity of 2.2 ± 0.3 molecules before addition of MMC (Fig 2A inset). This is clearly most consistent with a dimeric subunit of RecA in structures before MMC treatment. After MMC treatment, the most likely interval value is 3.1 ± 0.5 RecA-mGFP molecules, and estimating the additional unlabeled RecA content indicates a likely overall periodicity range of 3-4 RecA molecules (see Discussion and Figure S4).

In MMC-treated cultures we observe strikingly bright, elongated structures (Figures 3B-E). These resemble parallel or intertwined RecA-mGFP nucleoprotein filaments, that we denote as *bundles* following similar observations by others [21,28,34,42,54]. The bundles were identified in a pointillistic manner by overlaying the tracked foci with our measured localization precision of 40 nm. Though it is unclear whether this segmentation is able to distinguish individual filaments or bundles of RecA from one another, the segments reproduce the contiguous morphology of the bright structures at a diffraction-limited optical resolution (Figure 3C,F).

**Figure 3.**
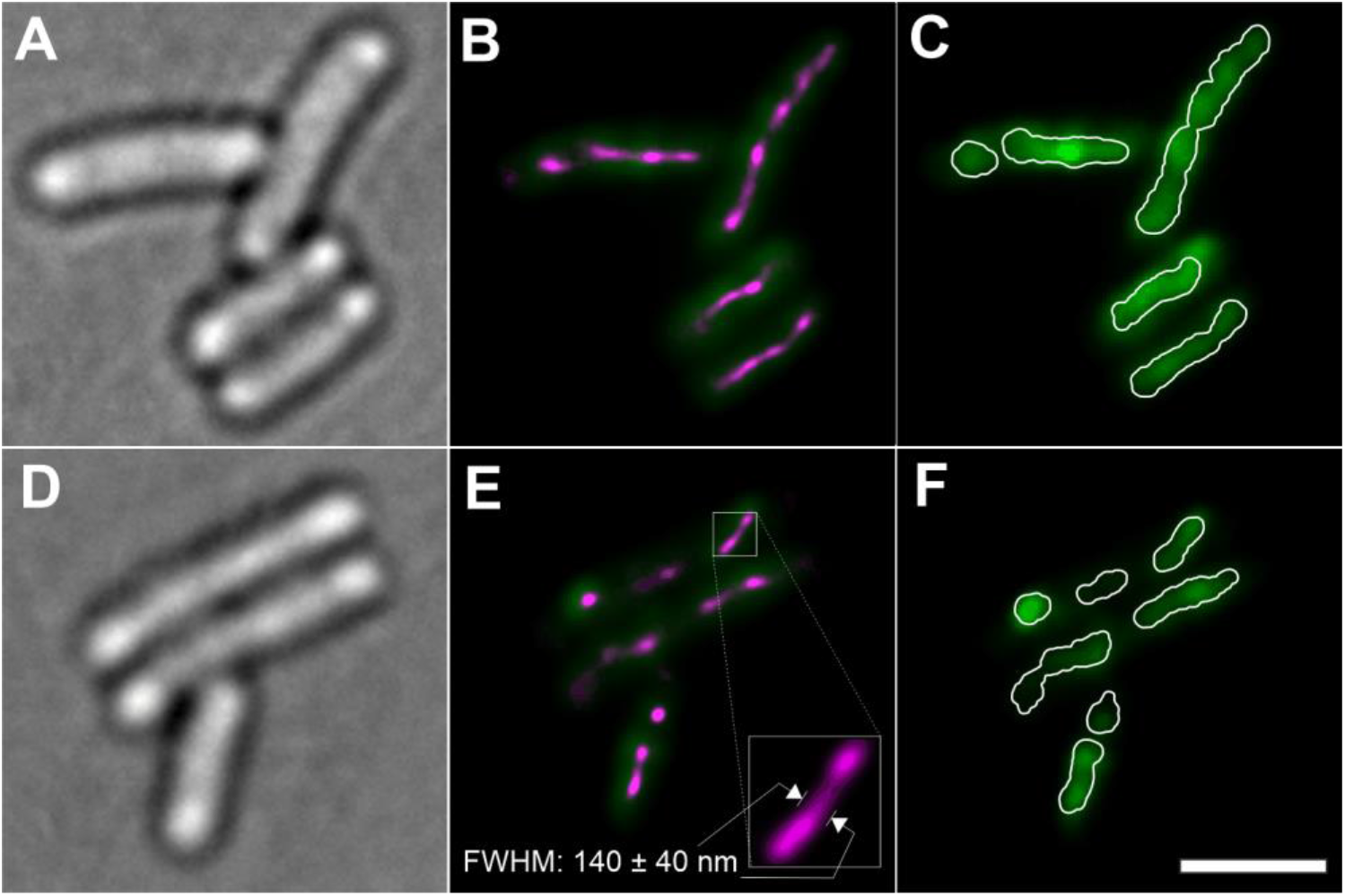
Bundles of RecA-mGFP filaments in MMC treated cells as observed in A,D) brightfield and B,E) The initial Slimfield fluorescent frame (green) overlaid with all super-resolved single-molecule tracks from the acquisition (*ca*. 40 nm spatial precision, with point localizations from foci visualized as a Normalised Gaussian rendering in ThunderSTORM, Materials and Methods 4.3.7), revealing filaments with high spatial precision (magenta); note that the contrast for the green Slimfield channel is set to half to aid the visibility of the superresolution rendering. C,F) Slimfield at full contrast, overlaid with segments derived from each super-resolved bundle by Otsu thresholding and expanding the resulting image masks by the point spread function width of 180 nm, so as to match the diffracted-limited widefield image optical resolution (white overlay); these segments were then used to calculate the segment protein copy number. Scale bar 2 μm.

Though single contiguous segments are evident along the full length of some cells (Figure 3C), the mean number of segments is 1.8 ± 0.4 per cell (Figure S3B). We occasionally observed several small segments per cell, in quantitative agreement (Figure S3B) with our expectation that segments occur at random under of a Poisson distribution, albeit conditioned on the presence of at least one segment persistently occurring per cell. Assuming that DNA damage also occurs randomly but under unconditional Poisson statistics, the number of unaffected cells can only be small when there is significantly more than one segment-inducing damage site per cell at any time. It is not clear how many DSBs per cell cycle an *E. coli* culture can sustain without loss of viability, but repeated stalling and collapse of the replisome is common, and cells with single chronic DSBs are known to replicate almost normally within the confines of an elevated SOS response [55]. Under the relatively mild MMC treatment conditions of our study, we observe a comparatively small fraction of the available RecA-mGFP in bundle-associated segments (here the sum of the segment protein copy numbers is <40% of the cell protein copy number, in contrast to 70% [34]). These observations favour an explanation that much of the RecA in elongated MMC-induced structures is bound to a relatively large number of ssDNA nicks as well as a small number of DSBs per cell at any one time.

The high estimated amount of RecA (Figure 2), and the substantial super-resolved breadth of these objects (Figure 3, Figure S3D) above the ~40 nm width of individual filaments [17] suggests that these are bundles comprised of either multiple RecA filaments, and/or multiple windings thereof. We calculate the segment protein copy number within each of these segments in a similar manner to each whole cell. We find the segment protein copy number is 2,800 ± 200 RecA-mGFP molecules (Figure S3A). That means each segment typically includes about three times as much RecA in total than the brightest polar assemblies detected in untreated cells (Figure S3A). Greater than 95% of these segments contain a track whose stoichiometry exceeds twice the mean stoichiometry of untreated cells. The RecA structures observed after MMC treatment cannot therefore be produced solely from the large RecA assemblies prior to MMC treatment, but most likely recruit additional RecA from the cytoplasmic pool. We cannot measure the ratio of RecA to available ssDNA directly, however, the high measured amount of RecA provides some indication that it occurs in high enough excess to form RecA-rich bundles rather than simple nucleoprotein filaments. The binding site density on each helical filament containing ssDNA was found in previous studies to be 1.5 nm per RecA in the presence of ATP [56,57]. As the individual filaments are known to be undersaturated with RecA under physiological conditions [58], one would expect a longer filament per molecule. In contrast, we find that each bundle-associated segment typically measures 900 ± 400 μm (mean ± s.d.) in length, 140 ± 40 μm wide (Figure S3C-E) and no greater than ~0.4 μm deep (based on depth of focus constraints), but contains a quantity of RecA we estimate sufficient to produce >7 μm total length of individual helical filament based on known structures [59]. The longest segments have a more efficient packing density of RecA-mGFP (Figure S3F) which approaches the binding site saturation limit of 1.5 nm / molecule. This link between length and efficiency could result from the functional alignment and elongation of the filament along the cell axis, meaning fewer re-entrant windings of any bundles, and exposure of vacant binding sites to free RecA in cytoplasm.

In contrast, the brightest RecA assemblies in untreated cells occur in isolation, and are never elongated but reside within diffraction-limited foci (Figure 1B). Defining these as containing RecA exceeding twice the mean labeled stoichiometry, these occur in 10 ± 3% of untreated cells and have a typical content of 800 ± 100 RecA molecules (Figure S3). This relatively high density is equivalent to >2 μm of filament packing inside a sphere <0.4 μm in diameter. While these assemblies resemble RecA storage bodies, as suggested previously [21], the *recA4155* R28A mutation has been shown to inhibit the formation of true DNA-independent storage bodies [24]. Despite the presence of wild type RecA, it is likely that our observations before MMC treatment indicate DNA-bound RecA bodies that are not filamentous.

The diffusive dynamics of RecA assemblies are also indicative of their state of condensation into filaments. Returning to the tracked foci of RecA-mGFP, we noticed that the mean diffusion coefficient decreases sharply from 0.17 ± 0.02 μm^2^/s to 0.07 ± 0.01 μm^2^/s following MMC treatment (Figure 4A). This initially low diffusivity, and the further drop in diffusivity, likely reflect the proportion of RecA condensed onto ssDNA. MMC induces formation of filaments and these are relatively static on the *ca*. 10 s timescale of the Slimfield acquisition. In contrast, we find that the mean diffusion coefficient of tracked RecB is not significantly affected by MMC treatment (Figure 4B), with untreated and treated values of 0.82 ± 0.03 μm^2^/s and 0.79 ± 0.03 μm^2^/s respectively (BM test, n=478, p=0.48 | NS). The diffusivity of RecB-sfGFP in tracks is still lower than expected for a single molecule freely diffusing in bacterial cytoplasm of ~10 μm^2^/s, based on simplistic assumptions of a hydrodynamic diameter of ~10 nm, and contrasts with the large amount of pool RecB-sfGFP that diffuse too quickly to be tracked. This observation hints at the tracked subset of RecB forming larger complexes with other partners not detected here, such as RecC and RecD.

**Figure 4.**
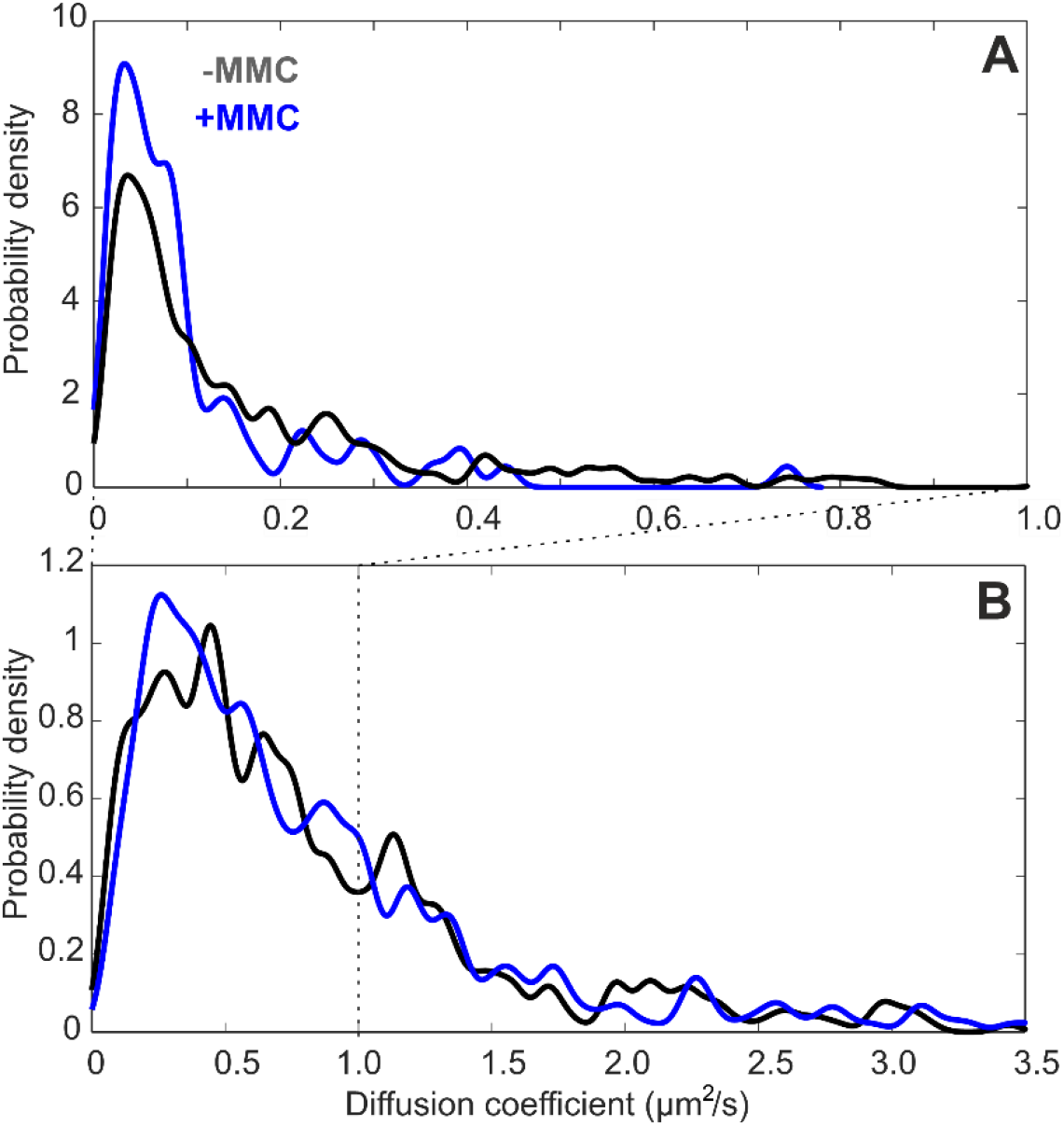
Distributions of instantaneous microscopic diffusion coefficient for tracks of A) RecA-mGFP and B) RecB-sfGFP obtained from Slimfield. Kernel density estimates were generated with a kernel width of 0.008 μm^2^/s corresponding to the lower bound uncertainty in diffusion coefficient, estimated as the localization precision / (timestep)^2^. Statistics are as shown for Figure 2.

## 3. Discussion

We used Slimfield to investigate the stoichiometry and spatial location of fluorescently tagged RecA and RecB proteins in live *E. coli* upon treatment with the DNA cross-linking and alkylating agent MMC. RecA and RecB are repair proteins whose involvement in MMC-specific damage repair pathways, as part of the SOS response or otherwise, is unclear. We probed the steady state effect of MMC on RecA and RecB at the minimum inhibitory concentration, which is relevant to sub-lethal antimicrobial exposure. Our results show that the sensitivity and dynamic range of Slimfield is sufficient to quantify counts, either by stepwise photobleaching of multi-molecular complexes or by direct detection of single molecules using millisecond sampling.

RecA assembly formation is not solely correlated with induced DNA damage. Before treatment with MMC, we find that a portion of RecA appears in foci at an average incidence of approximately 2 foci per cell. In 10% of cells, at least one of these foci is especially bright, circular and localized to one of the cell poles. A previous study reports that a minority of cells (4-9%) exhibit spontaneous RecA foci near the poles prior to DSB induction [34]. It has been suggested that wild type RecA foci at the cell membrane might act as nucleation points for later filament formation across DSBs [60], or that these are storage bodies outside the nucleoid [21]. However, the RecA-mGFP strain used here (and in [34]) is a *recA4155* (R28A) genotype which abolishes DNA-independent aggregation of RecA [52,61]. In this strain, we cannot eliminate the possibilities that wild-type RecA forms native storage bodies that are undetected due to exclusion of RecA-GFP, or indeed visible storage structures which do recruit the mutant RecA-mGFP (RecA4155), which would account for the resemblance of detected foci to previous observations of these bodies outside the nucleoid [21,34,61]. In the case where RecA-mGFP cannot participate in storage bodies and can only aggregate in the presence of DNA, there is an alternative explanation for the subset of RecA-mGFP foci we observe, distinct from membrane anchors and storage bodies. These foci do not appear to require RecB for spontaneous assembly [34] indicating that they are independent of DSBs and instead assembled at incidental sites of ssDNA. The foci lie consistently at the periphery of the cell, which indicates they are not likely to be associated with ssDNA within replication forks. These occasionally bright foci may instead simply reflect stochastic ssDNA nicks in a small proportion of cells of an otherwise healthy culture.

Our findings show that the RecA-mGFP copy number increases upon treatment with MMC. We observed a modest increase in the number of tracks, but whose stoichiometry per focus is almost twice those of untreated cultures. This observation of spatially localized RecA and is consistent with significant assembly formation ultimately leading to formation of long nucleoprotein filaments on ssDNA as nucleated from polar locations [34]. These filaments are known to accumulate into bundles as posited by Story et al. [42]. We observed filamentous bundles in MMC treated cultures, possibly due to increased availability of processed ssDNA from DNA damage sites. RecA-assisted homologous recombination and RecA* disassembly occur on a timespan between 15 min [17] and 2 hours [34]. We detect a large increase in RecA stoichiometry (Figure 2) and cellular protein copy number (Figure S5A) and decrease in diffusivity (Figure 4) even after 3 hours’ treatment, indicating that RecA bundles continue to form in response to constantly accumulating DNA damage.

Our observation of an about 2 intracellular segments per cell (Figure S3B) is consistent with approximately 2 MMC-induced RecA bundles each extending along opposite halves of a cell (Figure 3F) at any one time in the steady state. This observation may indicate the presence of a double-strand break (DSB) with nearly-bridged loci. However, according to the schemes in previous work [17,34], the development and breakdown of filaments [17] and bundles [34] takes typically <20 min, while for bundles only, recombination is the rate-limiting step, taking up to 90 min [34]. It follows that labeled bundles associated with DSBs would be expected to be bridged for most of their visible lifetime. It is therefore possible that either i) multiple DSBs are present and the segments correspond to different simultaneously bridged DSBs, or that ii) one bridged DSB is present alongside other defects which support RecA filament binding, such as ssDNA nicks.

Intracellular segments were typically aligned along the cell axis (Figure 3B,E) in agreement with the observations of filaments and bundles by other authors [17,34]. While some degree of alignment is expected for all segments much longer than the cell diameter (0.78 ± 0.05 μm), we note that more than half of the detected segments are shorter than this (Figure S3E), which may suggest an alignment mechanism that is not solely due to segement length. Moreover, segments appeared to follow the central axis of the cell, rather than the cell outline (Figure 3B,E), which suggests they fall mostly within the nucleoid rather than residing at the cell membrane, in keeping with the known DNA repair function of the filaments. Filament extension along the cell axis is not predicated on the presence of sister homology [34] but inherently reduces the dimensionality of the search for any homology to one across the cross-section of the cell, independent of cell length or DNA content [17]. Thus, extension vastly accelerates the search time [17]. However, the cause of the extension is unclear. It may reflect simple polymeric elongation under spatial confinement inside the cell, but extension is entropically unfavourable for a flexible polymer. Stiffening and/or thickening of filaments into bundles [34] would therefore faciliate extension. The bundle model in [34] suggested a thickened central backbone flanked by thin filament ends. The bundles observed in our study appear to be thickened with a typical cross-sectional full-width half-maximum of 140 ± 40 nm (Figure S3C) in agreement with previous observation, 160 ± 30 nm [34]. Rather than a monolithic central section, our observations resemble beads on a chain, or a sequence of thick and narrow sections (Figure 3B,E). We find the median width increases rapidly with segment length (Figure S3D), which in this binary framework, suggests the bulk of the increase in bundle length is taken up by the thickened portions and that the thin sections are relatively short. Yet, individual ~40 nm-wide filaments without thickened portions have also been observed previously to extend dynamically on the scale of minutes or less along the length of the cell [17]. We speculate that this suggests an active process of pole-to-pole translocation of thin filament ends (for example, as proposed in [59]), to facilitate the reduced search time.

The observable periodicity of RecA structures could indicate a difference in their macromolecular organization in response to MMC treatment. We observe a change in the periodicity of RecA stoichiometry from ~2 molecules in foci in untreated cells, to a ~3-4-mer within spatially extended filaments following treatment with MMC, after accounting for the unlabelled RecA content per cell with a correction factor of 1.3 ± 0.1 (Results 2.3 and Figure S4). Previous *in vitro* and *in vivo* studies indicate that RecA undergoes linear polymerization in a head-to-tail fashion, with dimeric nucleation points on ssDNA mediated by SSB [62] consistent with our finding of dimeric periodicity prior to treatment. These also provide evidence for stable trimeric, tetrameric, hexameric and the filamentous forms when ssDNA is present [63], consistent with our findings post-treatment. Our snapshot observation of filament stoichiometry cannot shed light directly on models of dynamic nucleation or stepwise growth, as explored in [64–66]. Rather, it explores molecular details of the characteristic protein subunits within the mature filament at steady state. The helical geometry of the filament, with a pitch of 6 RecA molecules per turn, implies that each group of 6 RecA forms a split-ring structure related to the intact hexameric ring of DNA helicases, but distorted axially such that rings each complete a single helical turn around ssDNA [67]. Such ring-shaped hexamers have been identified *in vitro* for both the wild type RecA protein, and the RecA (R28A) mutant [61] that is fused with GFP in our experiment. Even if isolated oligomers were somehow unstable *in vivo*, a polymeric filament could conceivably still result from a small, periodic barrier to polymerization corresponding to this split-ring distortion. This points to the hypothesis that the fundamental building block of RecA filaments is a factor of 6, if not a hexamer. However, our stoichiometry analysis suggests variability in the total size of assemblies, with our periodicity results indicating a range of 3-4 molecules per subunit. This could reflect trimers which form half-turns in the filament, or perhaps tetramers as an intermediate between preexisting dimers and hexameric rings. Although these data cannot directly establish whether independent oligomers of wild type RecA occur *in vivo* either on DNA or in the cytosol, it is conceivable that assembly and rearrangement of RecA subunits on DNA could generate the canonical ATP-inactive and ATP-active DNA-binding filaments [68,69]. In light of a recent study highlighting the role of RecN in RecA filament formation and activity [59], it is interesting to pose whether RecA assemblies with the dimeric subunit may be devoid of RecN and are ATP-inactive, and if these might then change to a higher oligomeric form upon DNA damage via the involvement of RecN and its associated ATP-activity.

Our measurements confirm that RecA has a very high concentration in the cytosol of live cells. We observe that untreated cultures comprise approximately 11,000 molecules of RecA-mGFP per cell, which increases to 20,000 RecA-mGFP molecules in cells treated with MMC. Of the latter, 28 ± 7% resides in filamentous bundles large enough to be resolved in millisecond widefield fluorescence images. Applying the approximate merodiploid correction factors that we estimated of 1.0 ± 0.1 and 1.3 ± 0.1 respectively (Results 2.3 and Figure S4), the total copy number is approximately 11,400 ± 200 RecA molecules in untreated cells, increasing to 25,300 ± 400 molecules in treated cells. Though less than the 4-5-fold transcriptional increase suggested by qPCR (Figure S4), the more than two-fold increase of total RecA with MMC resembles the increase detected in western blots (Figure S4). While the RecA copy number we estimate in untreated cells exceeds the *ca*. 5,000 molecules reported previously by Lesterlin et al [34], our more direct estimations are of similar order and correlate with previous work indicating 2,900-10,400 molecules, with the high end of this range obtained from cells in EZ rich medium using a ribosome profiling method [70]. Approximately 15,000 RecA molecules per cell in rich medium were reported previously, using semi-quantitative immunoblotting [71]; the same study found that the RecA copy number increased to 100,000 molecules upon MMC treatment. Large discrepancies between studies in the increase in RecA due to MMC treatment are not only due to treatment dose [72] but also arise from differences in *recA* genotype, culture media and growth conditions, as noted by others [21]. In particular, our study uses a minimally inhibitory treatment with MMC (Figure S1).

While the RecA-mGFP protein is not identical to native RecA in its enzymatic activity [32,43,52], the merodiploid strain used in our study also includes a full complement of native RecA from a single allele. This wild-type protein is expected to mitigate the partial loss of sensitivity in RecA-mGFP in any given mixed assembly, as shown on a cellular level by the similar SOS response profile [34] and lack of filamentation under treatment with MMC (Figure 1, Figure S1). The *recA* wild type allele is expressed under control of the native operator, while the *recA-mGFP* allele is expressed under the *recAo1403* operator. In the absence of treatment with MMC, this operator is known to result in an increased transcription rate of the *recA-gfp* allele relative to the wild type *recA* gene under its native promoter by a factor of 2-3, while both alleles are upregulated to the same level under induction of SOS [52]. Using quantitative Western blotting we estimate that prior to MMC treatment, RecA-mGFP is actually present at several tenfold more than the unlabeled protein (Figure S4), and that in the presence of MMC the ratio of RecA-mGFP to RecA is lower, at approximately 3-4 to 1. From these ratios, we derived our approximate correction factors of 1.0 or 1.3-fold for the total amount of RecA protein, in the absence or presence of MMC respectively. While RecA-mGFP is known to label RecA assemblies [34], it cannot form DNA-independent assemblies by itself [52], and is therefore reasonable to conclude that all labelled sites here represent occupied DNA on which wild type RecA and RecA-GFP are interchangeable. Even if the binding partition of wild type RecA were higher, for example reflecting the relative sensitivity (Figure S4A and [52]), the high relative concentration of RecA-mGFP (Figure S4B) would conceivably result in the majority of RecA sites on DNA being occupied by RecA-mGFP.

Lesterlin *et al*. showed that RecA immunostaining of filaments (agnostic as to GFP labelling) correlates with the fluorescent distribution of RecA-GFP [34], proving that dark filaments exclusively of wild-type RecA cannot be present. Though this result could potentially be interpreted that the structure is entirely RecA-GFP and that the highly sensitive wild-type RecA is excluded, this wild-type RecA would have to somehow rescue DNA repair function in the cytoplasm rather than in filaments, which has no known basis. We therefore assume the presence of hybrid filaments. In any case, the effect of excluding wild-type RecA from filaments would simply mitigate our periodicity correction factor toward unity, and narrow our estimate of the periodicity within filaments toward a value of 3 RecA molecules.

Unlike RecA-mGFP, we detected only modest quantities of RecB-sfGFP in untreated cells grown in minimal medium: 13.6 ± 0.5 molecules in tracks, and 126 ± 11 molecules in total per cell based on integrated GFP fluorescence corrected for cellular autofluorescence. Several previous reports also indicate that RecB is very scarce – typically less than 20 molecules per cell [35,73]. One of these studies estimated that there are just 4.9 ± 0.3 RecB molecules per cell using a HaloTag fusion allele labeled with HTL-TMR, and 4.5 ± 0.4 molecules per cell using magnetic activated cell sorting of the same RecB-sfGFP strain that we use here, albeit in M9 medium and restricted to nascent cells for which the average copy number is approximately halved [35]. An earlier mass spectrometry study used intensity based absolute quantification to estimate 9-20 RecB molecules per cell across different stages of growth in M9 minimal media [73]. Ribosome profiling estimated the RecB copy number to be 33-93 molecules per cell in different growth media [70]. However, these techniques are either *ex vivo* or necessitate significantly perturbed intracellular crowding that may conceivably result in potentially non-physiological molecular assemblies.

Comparing the number of RecB in tracks in our present study with the number of RecB in distinct foci per cell reported previously, we find a similar albeit slightly higher estimate, possibly because our approach is based on fluorescent fusions with a high labelling efficiency in unsynchronized cultures, as opposed to selecting nascent cells. However, our measurement of RecB copy number exceeds previous estimates. The large remainder in summed pixel fluorescence intensity may represent two possible contributions. The first is from RecB that diffuses faster than Slimfield can track. The highest diffusion coefficients of tracked RecB assemblies approach 3 μm^2^/s (95% quantiles, Figure 4). We estimate the limit of measurement as approximately 5 μm^2^/s, though it is conceivable that free monomeric RecB-sfGFP could exceed this, given that it has been estimated to reach diffusion coefficients equivalent to approximately 8 μm^2^/s in *E. coli* cytosol [74,75]. A second possible source is an increase in net autofluorescence relative to the parental strain when the real RecB are labeled; it is unlikely that could account for the discrepancy, since this would require a 3-fold increase in autofluorescence based on our measurements, and such a drastic increase lacks precedence (for example, upon treatment with MMC at a high level sufficient to induce widespread RecA filamentation, our estimation suggests only an increase in autofluorescence of no more than 20%). Furthermore, the measured rate of photobleaching of the diffuse RecB-sfGFP signal matches that of RecB-sfGFP tracks and is roughly half the rate of the autofluorescent parental cells (Figure S6, Table S2). The implication is that untracked RecB-sfGFP is the major contributor to mean cellular fluorescence, which is then a more accurate reflection of total copy of RecB than simply the number of molecules in tracks.

The cellular protein copy number of RecB does not change significantly with MMC (Table S1, Figure S5B,D), suggesting that there may only be a modest regulatory response to DNA damage. Although MMC is known to induce the SOS response and cell cycle arrest [4,48], *recB* expression is itself not induced directly as part of the SOS response. RecB-sfGFP foci increase neither in number (Figure 1J) nor stoichiometry (Figure 2B), which compares with earlier observations that treatment with MMC under similar concentrations to those used in our study do not significantly change RecB expression [22]. In fact, the number of observed tracks per cell *dropped* considerably after MMC treatment, due to a sharp increase in the proportion of cells in which RecB assemblies were absent, from 6% to 21%. This reduction in RecB assemblies was at odds with our expectation that MMC would eventually increase the recruitment of RecB in response to damage, if not increase the cellular production of RecB. MMC treatment is known to increase the occurrence of DSBs and thereby drive demand for DSB processing [4] that is typically mediated by RecB. Yet, rather than initiating cellular upregulation of RecB, treatment with MMC acts to partially deplete localized assemblies. Given that DSBs are likely to occur in the majority of cells under our MMC treatment, as indicated by the ubiquitous induction of RecA filaments (Figure 3), the fate of RecB assemblies cannot simply reflect the presence or absence of DSBs. The increase in the fraction of cells lacking RecB-sfGFP tracks is consistent with random, independent survival or breakdown of assemblies (Figure 1J). This result may indicate a situation where pre-existing RecB (hetero)complexes at foci are occasionally disassembled while interacting with sites of MMC-induced DNA damage, such as DSBs. This instability of (presumably heteromeric) RecB assemblies might result from successfully bridged pairs of RecA filaments, however, we did not detect any correlated loss of pairs of RecB foci, as might be expected for recombination events. Notably, the number of tracked foci detected per cell is approximately 2 for both RecA-mGFP and RecB-sfGFP. Future colocalization studies of RecA and RecB assemblies may offer more direct insight into the functional interaction and turnover of these repair proteins in regards to whether the average of 2 be related to the number of replication sites, or perhaps simply reflects a small average number of severe DNA damage sites per cell.

Independent of MMC treatment, we observed a dimeric periodicity for RecB-sfGFP. This suggests that RecBCD heterotrimers occur in pairs *in vivo*. Indeed, earlier *in vitro* studies identified the occurrence of (RecBCD)_2_ complexes, possibly held together by the nuclease domains of the two RecBCD monomers [12]. However, the authors concluded that the monomeric form is functional while the dimeric form is nonfunctional [13]. Furthermore, crystallization of the RecBCD complex for structural studies contained two RecBCD-DNA complexes in the asymmetric unit [76]. Our observations cannot determine covalent interactions directly between RecB molecules, but their cotracking is very strongly correlated. We can infer two details: first, that the dimeric form of the complex, (RecBCD)_2_, occurs in live cells, and second, that previous *in vitro* observations of dimers are carried over from their physiological state. Our findings suggest a hypothesis that assemblies with multiple pairs of RecB have a greater activity on DSBs than isolated RecB in the pool. Making the distinction between monomeric RecBCD in tracks and monomeric RecB in the untracked pool suggests that RecB monomers in the pool could potentially act as a reservoir. One may consider the alternative situation, where the monomeric pool are the functional RecB elements and the assemblies are reservoirs that disassemble in response to damage, but this makes less sense, since those monomers would already be in excess. A mean stoichiometry of ~6 molecules indicates that RecB foci may occur as colocalized assemblies that comprise roughly three pairs of RecBCD heterotrimers (Figure 5). It would be interesting to estimate the stoichiometries of RecC and RecD in future studies to understand their association in processing DSBs in greater detail.

**Figure 5.**
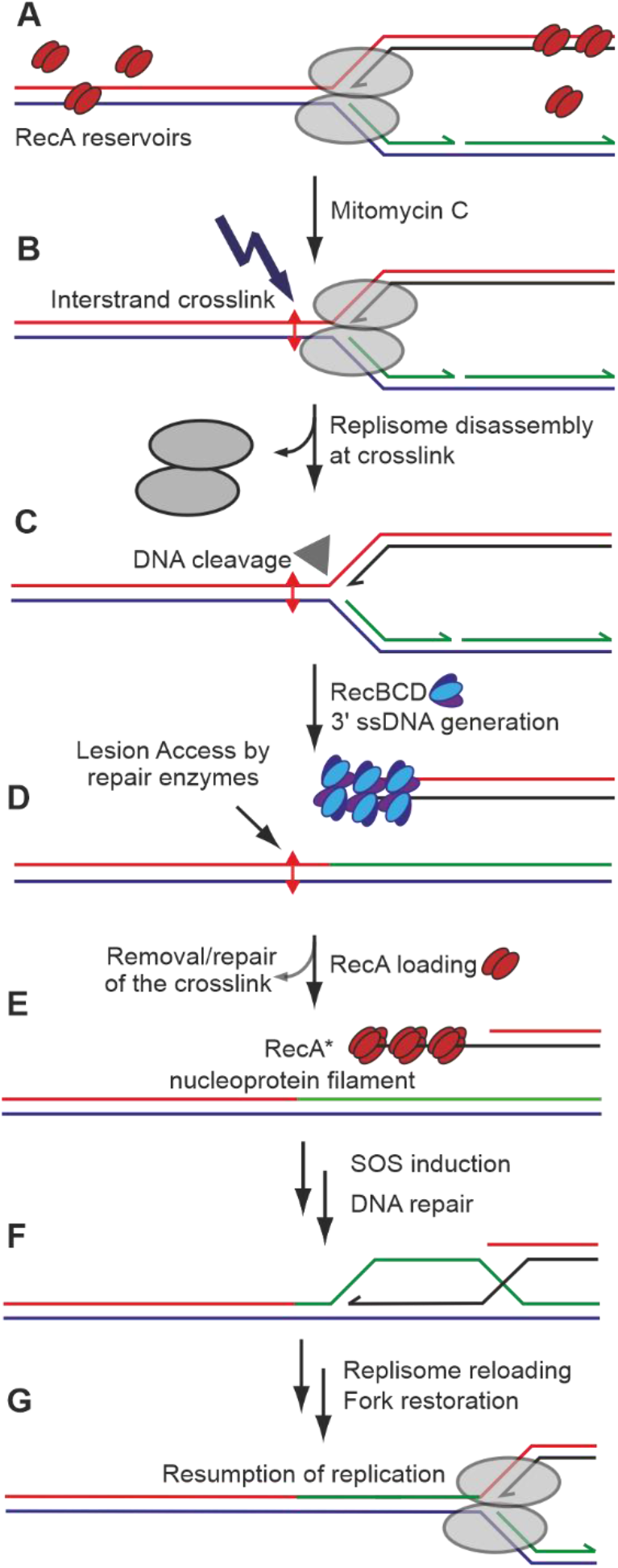
A model of DNA damage caused by treatment with MMC and subsequent repair at the replication site by RecA and RecBCD. A) Intact replication fork; occasional binding of multiple RecA dimers to DNA away from the fork as well as RecA dimers as DNA-free storage bodies in the cytoplasm; B) exposure to MMC and induction of an interstrand crosslink that acts as a barrier to an approaching replication fork; C) replisome dissociates if unable to overcome barrier; dissociated fork is recognised by branched DNA specific endonucleases (filled triangle) that can eventually cause DSBs leading to replication fork collapse; replication fork collapse allows access to repair enzymes to recognise the lesion; D) a newly generated DSB is recognised by RecBCD and processed to generate a 3’ single strand end; E) RecA dimers identify the newly generated ssDNA and assemble in groups of 3-4-mers into RecA* filaments; RecA is shown as a short stretch for illustrative purposes but may extend for many thousands of molecules over several hundreds of nm of ssDNA, and these filaments may be twisted and/or grouped into bundles. F) Strand exchange followed by processing of the double strand break, then recombination sufficiently upstream of the lesion and subsequent G) reloading of the replisome. This process allows sufficient time for the repair enzymes to repair the lesion on the template strand, so that replication may resume. For a detailed overview of the possible pathways to fork restoration, refer to [11].

While MMC-induced damage constitutes a range of chemical moieties [77], the canonical mechanism of MMC toxicity is of interstrand crosslinks at dG sites [6,7]. The specific repair of interstrand crosslinks (implied in Figure 5) can involve several repair pathways, primarily nucleotide excision repair (NER), which converts the crosslinks into dsDNA breaks [78]. Although NER enzymes such as UvrD typically degrade RecA filaments, NER is involved in the cleavage of damaged replication forks into suitable substrates for downstream processing, including RecA-mediated recombination [79]. Repair of the fork is then completed, for example by PriA-, Rep- and PriC-dependent pathways [11,18,80] on sets of ssDNA and a dsDNA end (Figure 5). The observation of a greater increase in RecA-mGFP copy numbers and foci compared to RecB-sfGFP could indicate a significant proportion of single-strand breaks and single-strand gaps at sites of crosslinks. While two previous studies reported that NER action on crosslinks also produces ssDNA nicks [78,81], we do not know if this applies strictly to MMC-induced NER, as our present work does not pertain to genes that *process* ss-gaps. Future analysis of proteins that process ssDNA breaks may potentially shed light on the relative occurrence of the two types of breaks by MMC and their relation to repair of replication forks.

While others have shown that *recB* deletion abolishes UV-induced filaments of RecA [34], we do not know the effect of *recB* deletion and MMC treatment on RecA dynamics. To avoid RecA interference in ‘normal’ ssDNA processes such as replication, the cell maintains strict control over filament nucleation, based on RecA and associated cofactor concentrations. It is therefore likely that the observed filamentation upon treatment with MMC is dependent on RecBCD, indirectly pointing towards increased occurrences of DSBs in these cells. Alternatively, if RecA nucleation is independent of RecBCD, one might anticipate little change in RecA dynamics upon *recB* deletion. However, further analysis of MMC-dependent RecA stoichiometry and copy number in a strain devoid of RecBCD activity - and with a controlled RecFOR pathway [19] – is needed to differentiate between these models.

In conclusion, RecA occurs as assemblies located near poles of wild type cells in a dimeric periodicity consistent with nucleation models. Upon mild treatment with MMC, RecA is upregulated at least two-fold, and assembles into long filamentous bundles on newly generated ssDNA in effectively all cells without exhausting the cytoplasmic reservoir. These mature bundles have a much lower diffusivity, reflecting their aggregation of a few thousand molecules each, with a structural periodicity in the range of 3-4 RecA molecules. The bundles are typically wider than single filaments, but both forms are known to facilitate homology search for homologous base-pairing with an intact duplex. Generation of ssDNA is known to occur at a DSB induced by processing of disassembled forks upon recognition by RecBCD. We observed RecB as a moderately diffusive set of three associated dimers at two locations in the cell, providing further evidence that RecBCD predominantly occurs as pairs of heterotrimers inside the cell at either end of DSBs. Our work implies the existence of a separate, significant reservoir of highly diffusive RecB monomers. Neither of these forms of RecB are upregulated upon MMC exposure, nor do they change their mobility. Accordingly, RecB is not a part of the SOS regulon. Instead, MMC-induced DNA damage impacts the formation – or induces a higher turnover – of these periodic RecB assemblies potentially associated with further DSB repair.

## 4. Materials and Methods

### 4.1. Strains, culture and MMC protocol

Three strains of *E. coli* were used in this study without alteration:

**Control:**

*MG1655*

**RecA-mGFP** [34]:

*MG1655 rpsL (Str*^*R*^,*lac+) ygaD1::kan recAo1403 recA4155,4136-gfp901, fhuB::recAwt-cm*

**RecB-sfGFP** = MEK706 [35]:

*MG1655 recB::sfGFP*

The RecA-mGFP strain used here is the same as in [34]. It is a merodiploid, natively promoted derivative of the SS3085 strain [52,82,83]. It expresses both i) the wild type unlabelled RecA protein from a single, ectopic wild-type *recA* allele, under a wild-type operator, and ii) a labeled mutant protein at the native *recA* site with a mutant operator *recAo1403*, which has a higher transcription rate than wild type [52]. The labelled *recA* is a *recA4155* (R28A) mutant, which complements the wild type recombination function *in vitro* [34,61] but has important differences in self-assembly. Unlike wild type RecA, the *recA4155* forms of RecA cannot alone form assemblies independently of DNA as proven by comparisons *in vitro* [61] and competitive binding studies *in vivo* [52]. The *gfp-901* label is the same as *mut2* (A206T) which corresponds to a monomeric GFP (mGFP) [84]. The notation *recA4136* refers to the insertion of a linker, as well as the *gfp-901* gene, between the penultimate and ultimate stop codons of *recA*. The fusion with mGFP impairs the recombinant sensitivity of the labeled RecA protein; were the strain to include only the fused allele, it would be fully as SOS inducible as wild type MG1655, but only approximately two-thirds as UV resistant, and would be compromised up to *ca*. 10-fold for recombination activity [21,52]. Induction of the SOS response would also take 30 min, or roughly twice as long as wild type (the R28A mutation prevents this from being an additional 2× slower) [52]. The merodiploid strain rescues both these functions and their kinetics: functional RecA filaments labeled with 70% of the total available mutant fusion protein form within just 15 min of DNA damage and reach steady state within 90 min, similar to wild type MG1655 [34].

The RecB-sfGFP fusion was constructed in [35] by wild-type *recB* replacement under plasmid-mediated recombination. N-terminal fusions were shown in [35] to be functional using growth curves and tests of DNA repair, in contrast to C-terminal fusions which may disrupt RecBCD complexation [35].

*E. coli* strains were grown overnight in 56-salts minimal media at 30°C to mid-log phase in an Innova 44 shaker incubator (New Brunswick). The mid-log phase cultures were concentrated to ~100 cells/ml (OD_600_ ~0.3) and split into two equal fractions. Aliquots were adjusted to either nil (MMC-) or the minimum wt inhibitory concentration of 0.5 μg/ml MMC (MMC+) (Figure S6 and [48]) and incubated at 30°C for a further 3 h (Figure S6). Cells were harvested for microscopy on 1% w/v agarose pads suffused with the same liquid media and imaged within 1 h.

### 4.2. Slimfield

A custom-built Slimfield microscope was used for single colour, single-molecule-sensitive imaging with a bespoke GFP/mCherry emission channel splitter as described previously [11,37]. The GFP channel was recorded, while the mCherry channel was used only as a negative control. The setup included a high-magnification objective (NA 1.49 Apo TIRF 100× oil immersion, Nikon) and the detector was a Prime95B sCMOS camera (Photometrics) operating in 12-bit gain at 180 Hz and 3 ms exposure/frame, for a total magnification of 53 nm/pixel. The samples were illuminated either in brightfield, or for Slimfield fluorescence in camera-triggered frames by a collimated 488 nm wavelength continuous wave OPSL laser (Coherent, Obis LS) in Gaussian TEM_00_ mode at a power density of 5 kW/cm^2^. The number of frames per acquisition was 2,000 for RecA and 300 for RecB strains.

### 4.3 Quantitative tracking and protein copy number analysis

#### 4.3.1 Identification of Slimfield foci and assignment into tracks

Slimfield image sequences were processed by custom ADEMscode software in MATLAB (Mathworks) [33,80,85–87]. This pipeline identified foci from local maxima in pixel values within individual frames. An iterative Gaussian mask algorithm was used to detect the centroids of foci, using a circular region of interest of radius 5 pixels within a sliding window of 17 pixels. The intensity of each focus was calculated as the sum of the circular region corrected for the average background in the surrounding annular region. The prospective foci were accepted if their intensity was >0.4× the standard deviation in the background region. The nearest neighboring foci in adjacent frames within 8 pixels of each other were assigned to the same track, with a minimum of 4 foci per track. The typical track duration was limited by diffusion and/or photobleaching to a mean of >13 foci per track over ~75 ms real time, or ~40 ms cumulative exposure (Table S1).

#### 4.3.2. Diffusion coefficient

The centroids of the foci within each track, as generated from the ADEMScode tracking analysis above, were used to calculate displacements over the length of each track in chronological sequence. From these, the mean square displacements (MSDs) of each track were calculated by averaging the square of the displacements across equal lag times, corresponding to all possible intervals between frames up to the length of the track. For each track, the MSDs at the four lowest lag times were linearly interpolated (with a constraint on the fit of passing through a specified intercept on the lag time axis, equal to the square of the measured localization precision of 40 nm divided by the frame interval of 5.7 ms). The initial slope of this fit (and corresponding error) was then divided by a factor of 4 according to the 2D diffusion equation [88] to yield a diffusion coefficient (and error) for that track.

#### 4.3.3 Characteristic single-molecule brightness

The intensity of each focus was estimated by integrating the local pixel values with a local sliding window background subtraction. After photobleaching sufficiently to show single photoactive GFP molecules, the characteristic single-molecule brightness of a single GFP molecule was estimated from the modal brightness of these foci. These were confirmed to be broadly consistent with estimates of the signal per GFP in each dataset were determined from the monomeric intervals in total number of counts due to stepwise photobleaching, as identified by a Chung-Kennedy edge-preserving filter (15 ms window, 50% weighting, Figure S7) [89]. This integrated intensity is characteristic for each fluorescent protein under fixed imaging conditions, although mGFP and sfGFP were found to be indistinguishable in this respect, and hereafter referred to collectively as GFP. To ensure consistent counts per single-molecule probe, analysis was restricted to the uniformly illuminated area lying within half of the 1/e^2^ beamwaist of the excitation laser in the sample plane. The integrated intensity of GFP *in vivo* was found to be within 14% and 9% respective errors in RecA and RecB (88 ± 18 and 177 ± 16 pixel grey values per GFP for the respective gain modes). The combined equivalent is 88 ± 7 photoelectrons per GFP per frame, which is precise enough to unequivocally identify groups or steps of up to 12 GFP molecules.

#### 4.3.4 Stoichiometry

Each track is associated with an assembly that contains a certain number of molecules, or stoichiometry, at the initial point of acquisition. To estimate this stoichiometry for a given track, the intensities of the constituent foci were linearly extrapolated using the first 4 datapoints in the track back to the timepoint of initial laser exposure. This initial intensity of this fit was divided by the characteristic single-molecule brightness signal associated with one fluorescent protein under a fixed excitation-detection protocol. The result is a stoichometry expressed as a number of molecules. The standard error associated with a stoichiometry value of 1 molecule is approximately 0.7 molecules. To avoid undercounting bias due to photobleaching, only tracks in the first 10 frames after laser exposure were considered for stoichiometry estimates.

#### 4.3.5 Periodicity

The distributions of track stoichiometry may show periodic peaks, whose smallest reproducible interval can be interpreted as a physical repeat unit or *periodicity* within assemblies. To calculate periodicity, first the stoichiometries of all tracks within each acquisition were represented as a kernel density distribution. The kernel width used was the empirical standard deviation on the characteristic single molecule brightness of 0.7 molecules [41]. Peaks in this distribution were detected using the MATLAB *findpeaks* function, and the intervals between nearest neighbor peaks were calculated. These sets of nearest neighbor intervals for each acquisition were then aggregated across the relevant population of cells. A second kernel density estimate was calculated over the intervals for a population, with a kernel width of 0.7 molecules multiplied by the square root of the mean stoichiometry, divided by the square root of the number of interpolated intervals. The fundamental value of this interval distribution (corresponding to the center of the leftmost peak in Figure 2 insets) was refined by fitting the curve with a sum of Gaussian terms centerd at multiples of the fundamental value. To accommodate the uncertainty in the single molecule characteristic brightness, the fundamental value of the fit was not constrained to an exact integer value but represents a heuristic model for the periodicity. The number of terms in the fit was set to minimize the reduced χ^2^ metric in the fit. This modal value was reported with 95% confidence interval as the periodicity of assemblies in each population. This method of estimating periodicity was verified as independent of the mean stoichiometry using simulated data drawn from noisy Poisson-distributed multiples of an oligomeric ground truth (artificial input value). This analysis reproduced the expectation that the minimum number of tracks required for sufficient peak sampling, and therefore the limit of periodicity detection, scales with the square root of the mean stoichiometry.

#### 4.3.6 Cellular protein copy numbers and pool stoichiometry

The cellular protein copy numbers as reported in the Results Section 2.1, Table S1, Figure S5 and Discussion correspond to whole cell masks, as identified using the manual annotated machine learning segmentation output from brightfield images (Figure S2 and Supplementary Methods). Integrated intensities of cells (uncorrected cellular protein copy numbers) and pool stoichiometries, were determined not from tracked foci, but directly from the raw image sequences using the CoPro package in ADEMscode software following [37] with the characteristic single-molecule brightness of GFP (as described in 4.3.2), the cell masks, and the camera’s dark pixel bias as input. The procedure effectively adds up all of the pixel values within the mask in question in an initial frame, and accounts for the convolution of the 3D cell volume with the widefield point spread function, followed by projection onto a 2D image. To obtain the cellular protein copy number in the labeled strains, and account for the contribution of autofluorescence, we calculated the difference in mean integrated intensity per segment between the labeled and parent strains under the corresponding MMC± condition, adjusted by the ratio of mean segment area. The pool stoichiometry in each cell is a measure of its untracked molecular concentration. It is calculated in CoPro as the cell’s integrated intensity, less the mean integrated intensity of the parental cells, less the total stoichiometry of tracked foci in the cell, divided by the area of the cell mask relative to the area within one diffraction limited focus.

#### 4.3.7 Super-resolved images and segment protein copy numbers of RecA-mGFP

The segment protein copy numbers as reported in Results Section 2.3 and Figure S3, were calculated with ImageJ, using as input the segments corresponding to bundles or foci, instead of whole cells. These segments were obtained starting from the coordinates of localized, tracked foci in Slimfield analysis, from the latter stages of photobleaching below a threshold stoichiometry of 2 molecules; these were imported into ThunderSTORM software [90]. The Visualization module to build a pointillistic super-resolved image at 40 nm lateral spatial precision (as shown in Figures 3B,E) at 5× upscaling (11 nm pixel size), which was then smoothed with a Gaussian filter of 4 pixels’ width, and automatically Otsu thresholded to generate a superresolved binary mask. The masks were then expanded by a distance equal to the widefield resolution of 17 pixels (~180 nm) to match the features in the Slimfield images. The integrated intensities were extracted, as in [91], from the sum of fluorescent pixel counts in the Slimfield images (Analyze Particles > Multi-Measure function in ImageJ) less the area multiplied by the camera pixel dark value. To yield segment protein copy numbers (Figure S3A), the resulting integrated intensities were corrected for the relative autofluorescence, by subtracting the integrated intensity of parental cells adjusted by by the ratio of mean segment area. The Multi-Measure output also included the Feret diameter of each segment which was used as an estimate of its end-to-end length (Figure 3D-F).

#### 4.3.8 Photobleaching rates

Photobleaching rates were estimated by fitting the decrease in background-subtracted cellular protein copy number or mean track stoichiometry over the exposure time using MATLAB *cftool*. The fit consisted of a monoexponential decay to the first 10 frames with variable initial intensity and decay constant, but with a baseline fixed to the average intensity after 50 frames. Fits were then refined to include only data within the initial 1/*e* decay time (Table S2). RecA-mGFP and RecB-sfGFP photobleach decay times were consistently dissimilar at 13 ± 2 and 6 ± 1 frames respectively; sfGFP is typically several-fold less photostable than comparable enhanced GFPs under high intensity illumination [92].

#### 4.3.9 Statistical tests

We performed multiple statistical comparisons on each set of tracked data (typically ~5: number of tracks, stoichiometry, periodicity, diffusivity, copy number), which we account for using the standard Bonferroni correction; the significance level is adjusted downwards by a factor of the number of comparisons, α = 0.05/5 = 0.01).

### 4.4 Gene expression assays

#### 4.4.1 Quantitative PCR (qPCR)

Treated and untreated cultures were grown as in section 4.1. Total RNA was then isolated using Monarch Total RNA Miniprep Kit (New England Biolabs). cDNA was synthesised from 350 ng of total RNA from each sample using Superscript IV reverse transcriptase (Invitrogen) according to the manufacturer’s instructions using random hexamer primer (ThermoScientific).

The cDNA was then subjected to qPCR using Fast SYBR Green Master Mix (ThermoFisher) in a QuantStudio 3 Real-Time PCR System. The *recA* primer pair amplified *recA* cDNA in the wild type and both *recA* and *recA-GFP* mRNA in the labeled strain. *recA-GFP* alone in the labeled strain was amplified using *GFP* primer pair. 16s rRNA was used as a housekeeping control.

Data obtained was analysed using the standard curve method [51]. Standard curves were generated from serial dilutions of PCR products with known concentrations derived from genomic DNA. Fold increase in mRNA levels was calculated by dividing the values obtained for treated mRNA with the untreated. Results are shown in Figure S4A.

#### 4.4.2 Western blots

Six samples of normalised *E. coli* cell cultures were prepared as above (section 4.1) in 1 ml aliquots at OD_600_ ~0.2. The cells were isolated using centrifugation at 10,000 × *g* for 2 min in a microfuge to prepare them for SDS-PAGE / immuno-detection. The cell pellets were resuspended in 75 μl of SDS loading buffer and boiled for 5 min at 95°C before application of 15 μl onto a 4-20 % gradient gel. The gel was subsequently transferred to nitrocellulose and the membrane was placed in blocking solution (PBS-T, 5% (w/v) non-fat milk) for 3 h. Primary antibody (anti-RecA) was incubated at 1/500 overnight in blocking solution before the membrane was washed (4 × 5 min) in blocking solution. Secondary antibody (goat anti rabbit-HRP) was incubated at 1/2,000 dilution for 4 h in blocking solution before the membrane was again washed (4 × 5 min) in blocking solution. A final wash in PBS was performed before development using ECL and image acquisition (iBRIGHT). Results are shown in Figure S4B.

## Author Contributions

A.P-D, A.S, and M.L. designed the research; A.S. cultured and treated cells; A.P-D performed microscopy and data analysis, visualization and curation, J.S. and L.F. wrote and validated segmentation software; A.P-D and A.S drafted the paper; A.P-D, A.S, J.S. and M.L. edited the paper; M.L. supervised and administered the project. This research was funded by BBSRC, grant numbers BB/P000746/1 and BB/N006453/1, and EPSRC, grant number EP/T002166/1.

Data Availability Statement: The raw imaging data is available on reasonable from https://doi.org/10.5281/zenodo.6639101; the MATLAB tracking analysis code can be found at https://github.com/alex-payne-dwyer/single-molecule-tools-alpd. The U-Net image segmentation architecture originated from code obtained from the NEUBIAS Academy workshop (http://eubias.org/NEUBIAS/training-schools/neubias-academy-home/).

## Acknowledgments

The authors thank Dr. Christian Lesterlin for the gift of the RecA-mGFP strain and Prof. Meriem El Karoui for the RecB-sfGFP strain. We thank the Biosciences Technology Facility for technical assistance with gene expression assays.

