## Supplementary methods for "RecA and RecB: probing complexes of DNA repair proteins with mitomycin C in live *Escherichia coli* with single-molecule sensitivity"

**Supplementary tables**

| | RecA-mGFP (≤ total RecA) | | RecB-sfGFP | |
|---|---|---|---|---|
| *MMC treatment* | - | + | - | + |
| *Number of cells* | 190 | 67 | 249 | 307 |
| *of which contain tracks* | 170 | 60 | 234 | 242 |
| *Projected cell area (μm²)* | 1.8 ± 0.2 | 2.1 ± 0.2 | 1.5 ± 0.1 | 1.6 ± 0.1 |
| *Number of tracks* | 316 | 125 | 514 | 478 |
| *Number of tracks per cell* | 1.66 ± 0.06 | 1.86 ± 0.16 | 2.06 ± 0.09 | 1.56 ± 0.06 |
| *Number of tracks per cell containing at least one track* | 1.86 ± 0.07 | 2.08 ± 0.17 | 2.20 ± 0.07 | 1.98 ± 0.07 |
| *Integrated intensity per cell (equivalent GFP molecules)* | 11,500 ± 200 | 19,600 ± 300 | 185 ± 7 | 174 ± 6 |
| *Autofluorescent integrated intensity per cell (equivalent GFP molecules)* | 71 ± 9 | 97 ± 17 | 59 ± 9 | 73 ± 13 |
| *Cellular protein copy number (molecules)* | 11,400 ± 200 | 19,500 ± 300 | 126 ± 11 | 101 ± 14 |
| *Sum of stoichiometries per cell (molecules)* | 510 ± 30 | 1080 ± 60 | 13.6 ± 0.5 | 9.5 ± 0.3 |
| *Sum of stoichiometries per cell with tracks (molecules)* | 570 ± 30 | 1210 ± 70 | 14.4 ± 0.5 | 12.1 ± 0.3 |
| *Stoichiometry (molecules)* | 310 ± 8 | 580 ± 30 | 6.59 ± 0.14 | 6.10 ± 0.19 |
| *Autofluorescent pool stoichiometry (molecules)* | 0.23 ± 0.03 | 0.27 ± 0.04 | 0.23 ± 0.03 | 0.27 ± 0.04 |
| *Pool stoichiometry (molecules)* | 33.6 ± 4.3 | 46.0 ± 3.2 | 0.46 ± 0.07 | 0.35 ± 0.10 |
| *Diffusion coefficient (μm²/s)* | 0.17 ± 0.02 | 0.07 ± 0.01 | 0.82 ± 0.03 | 0.79 ± 0.03 |
| *Track duration, cumulative exposure (ms)* | 49 ± 3 | 111 ± 9 | 39 ± 3 | 40 ± 3 |
| *Track duration, real time (ms)* | 93 ± 4 | 212 ± 21 | 74 ± 5 | 76 ± 4 |

**Table S1.** Average properties of molecular tracks for each strain and condition. Values shown as mean ± SEM. Amounts of RecA shown represent only direct measurements of RecA-mGFP and do not account for unlabeled RecA. The autofluorescent metrics (integrated intensity per cell and pool stoichiometry) are the values found for the MG1655 control strain, adjusted by the ratio of mean cell area of the labeled strain to that of the control strain in each condition. The *track duration* refers to the total exposure time for which that track was detectable above noise, for example before leaving the detection volume or photobleaching entirely; above an intrinsic limit due to the optical detection efficiency and photophysics of the fluorophore, it becomes an indicator of the brightness and immobility of the detected objects, in this case the condensed state of RecA-mGFP after MMC treatment.

| **Signal component** | **Photobleach decay time (frames)** |
|---|---|
| *Autofluorescence* | 3.1 ± 0.8 |
| *RecA-mGFP* | 13.9 ± 1.7 |
| *RecA-mGFP in tracks* | 12.3 ± 1.8 |
| *RecB-sfGFP* | 6.2 ± 1.1 |
| *RecB-sfGFP in tracks* | 6.0 ± 0.9 |

**Table S2**. Characteristic photobleaching decay times (mean ± SEM) as determined from refined monoexponential fits in Figure S6 (see caption of Figure S6 for associated numbers of cells and foci).

**Supplementary figures**

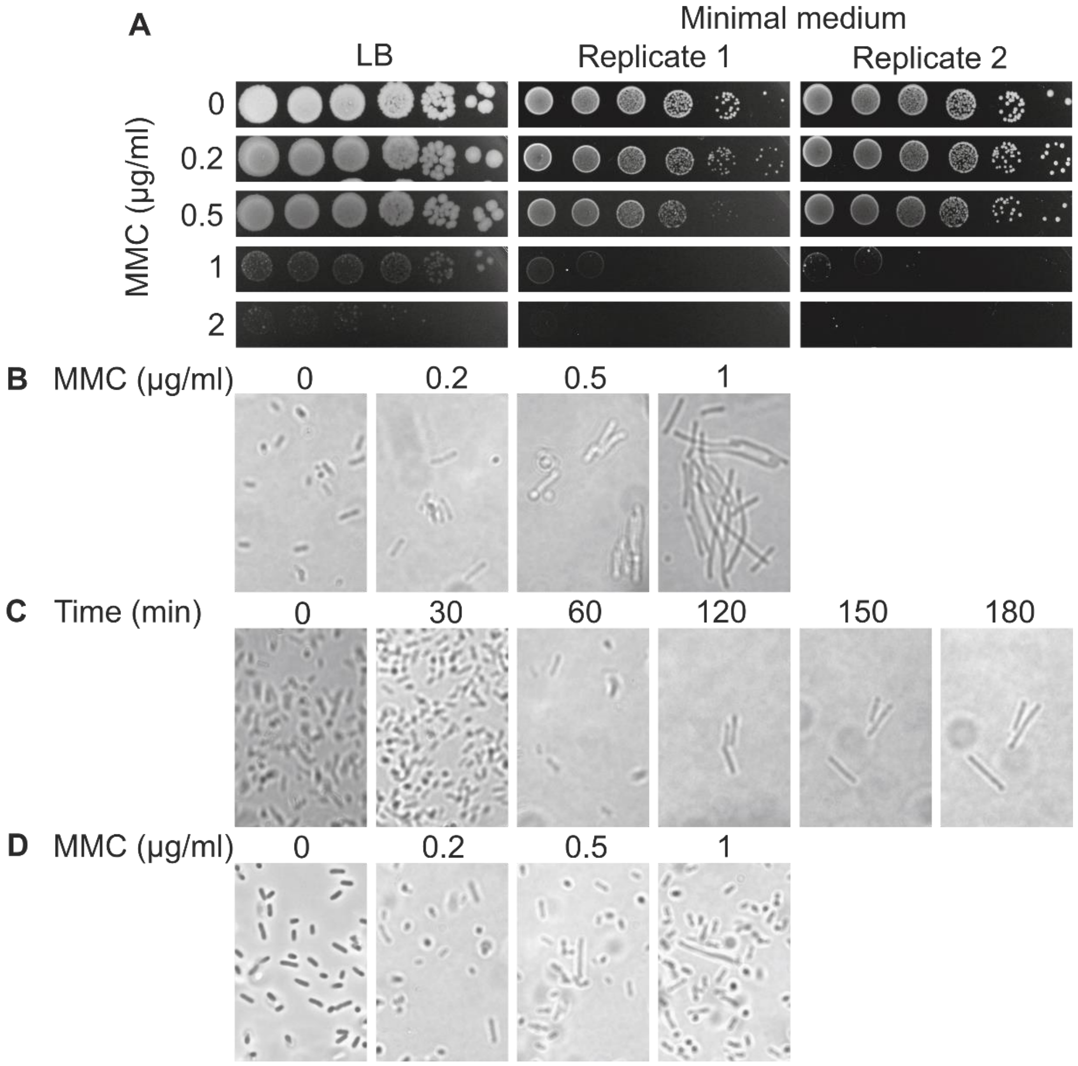


**Figure S1.** Determination of the optimal mitomycin C concentration and treatment time for Slimfield microscopy. A) Serial ten-fold dilutions of wild type *E. coli* MG1655 cells were spotted on a series of LB plates at various concentrations of mitomycin C and incubated overnight at 37°C to determine the maximum concentration of mitomycin C at which cells are viable. The results indicated that 0.5 µg/ml was the maximum concentration at which *E. coli* cells were viable under our rich-media growth conditions. B) Cells were grown in minimal medium to mid-log phase at various concentrations of mitomycin C and observed under the microscope to determine the concentration at which minimal filamentation occurs. A filamented cell phenotype is indicative of DNA damage, however excessive elongation of cells complicates quantitative high magnification microscopy and exacerbates the variance across clonal cells of different drug susceptibility. The results indicated that at 0.5 µg/ml mitomycin C there was minimal filamentation of cells so was conducive to microscopy. C) Mid-log phase cells grown in minimal medium were exposed to 0.2 µg/ml mitomycin C for varying amounts of time to determine the time at which excessive filamentation occurs. Note that the apparent changes in cell number density reflect unrelated variations in initial cell seeding, rather than survival over time. D) Based on the results in B and C, mid-log phase cells, this time grown in minimal medium, were exposed to various concentrations of mitomycin C for 180 min. It was concluded that 0.5 µg/ml mitomycin C for 180 min is an appropriate combination of maximal concentration and maximal exposure duration at which cells exhibit minimal filamentation indicative of DNA damage without compromising viability or morphology.

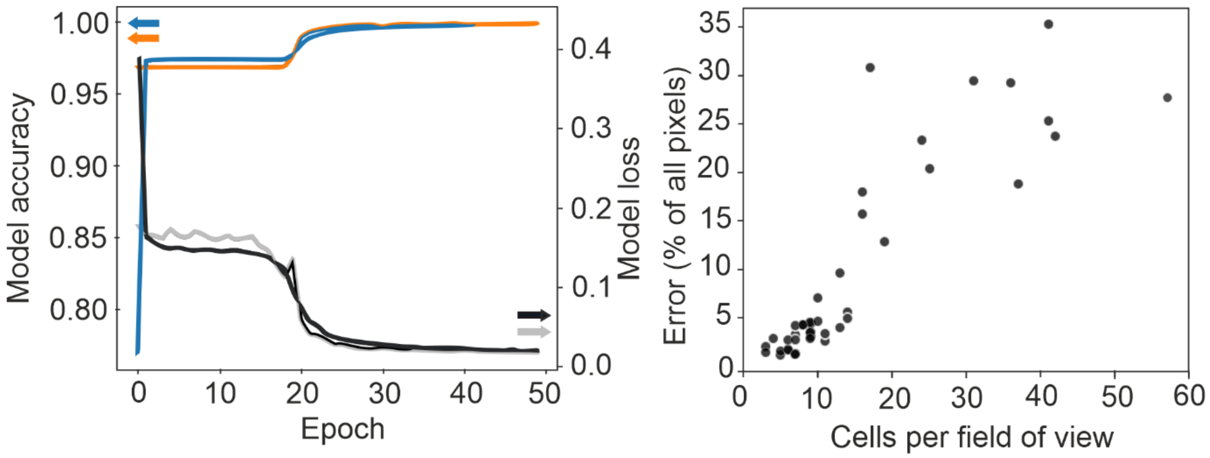


**Figure S2.** A) The accuracy and loss of the machine learning model for cellular segmentation (Supplementary Methods), as a function of epoch (a machine learning term referring to the training iteration) for the training images (blue, black respectively) and validation images (orange, grey respectively) using the U-Net with optimal parameters; B) rate of classification error for pixels in image against number of cells per field of view for all experimental datasets.

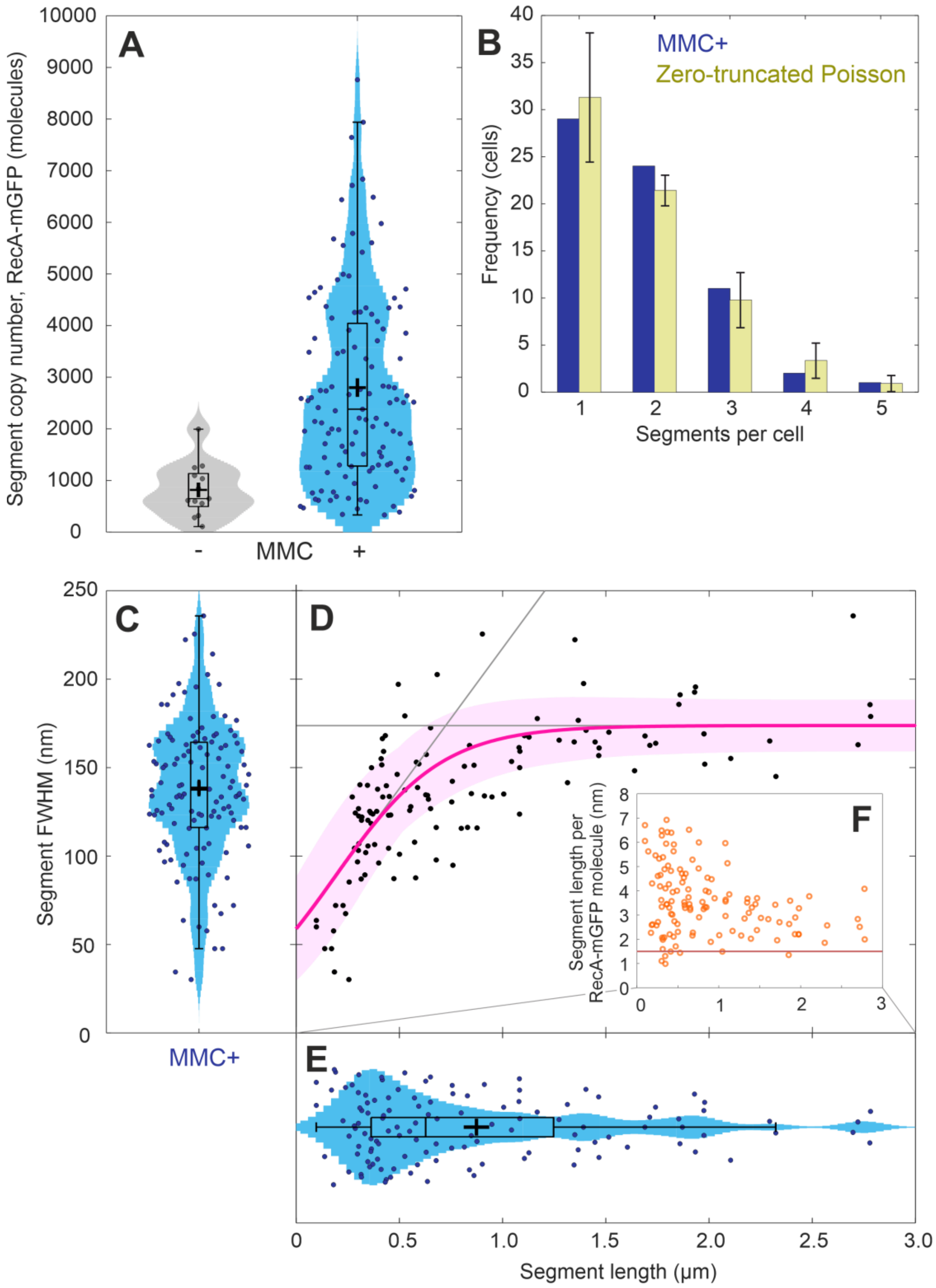


**Figure S3.** Size, incidence and spatial extent of super-resolved segments containing RecA-mGFP based on Figure 3B,E (prior to the widefield expansion of segments shown in Figure 3C,F). Each subcellular segment was determined by thresholding the superresolved images rendered using ThunderSTORM software [89] from precisely localized Slimfield tracks in untreated cells (grey), identified with prefilamentous RecA assemblies (n=13 segments in N=190 cells), and in MMC treated cells (blue), identified with active RecA bundles (n=123 segments in N=67 cells) as shown in Figure 3. On MMC treatment, A) not only does the number of segmented RecA-mGFP objects increase, but also the amount of RecA-mGFP in each increases by several fold, reflecting the drive toward filamental condensation. B) the number of segments per cell associated with RecA bundles (blue columns) closely follows a zero-truncated (conditional) Poisson distribution (yellow) with errors corresponding to 99%

CI on a maximum likelihood fit in MATLAB's *cftool*, yielding an corresponding rate parameter of $\lambda = 1.3 \pm 0.3$ segments per cell, but a mean of $1.8 \pm 0.4$ segments per cell. This matches the empirical mean of $1.8 \pm 0.1$ segments per cell. All cells detected contained at least one RecA-mGFP segment, which is not an observation consistent with a standard Poisson distribution of any mean in the feasible range. The superresolved full-width half-maximum (FWHM) of RecA-mGFP segments in treated cells C) is extracted by taking three randomly sampled cross-section profiles in each segment on the superresolved image (ImageJ line profile), estimating the FWHM of each, and the segment FWHM is reported as the median of these three FWHMs. This analysis reveals that FWHM of a segment is D) an increasing function of its end-to-end length; the line of fit (magenta) is an empirical sigmoid function of the form $f(x)=1/(exp(-ax+b)+c)$; applied in MATLAB's *cftool* (3 degrees of freedom, Pearson's $R^2$ = 0.55), with 95% CI bounds (magenta area). The range of lengths E) extends up to 3 µm, reflecting the maximal dimensions of the dividing host cell in the precursory stages of filamentation. The FWHM ceases to increase for segments much longer than ~0.7 µm, which coincides with a typical cell diameter. The limit in the FWHM line of fit is 170 ± 15 nm (95% CI), which is not statistically greater than the mean FWHM of all segments, 140 ± 40 nm. The intercept is 55 ± 20 nm (95% CI) and the initial slope is 0.14 ± 0.05 (95% CI) denoted by gray lines. The packing density of RecA-mGFP molecules is well represented by F) the effective distance per monomer along the length of the segment (approximated here as the segment copy number divided by the end-to-end length). Longer segments have a more efficient packing which approaches the limit of a fully-consolidated single filament of RecA on ssDNA (1.5 nm / molecule) [55]. Note these estimates of cellular protein copy number correspond only to labeled RecA-mGFP; they may be corrected by a factor of 1.3 ± 0.1 to account for unlabeled RecA in the merodiploid strain.

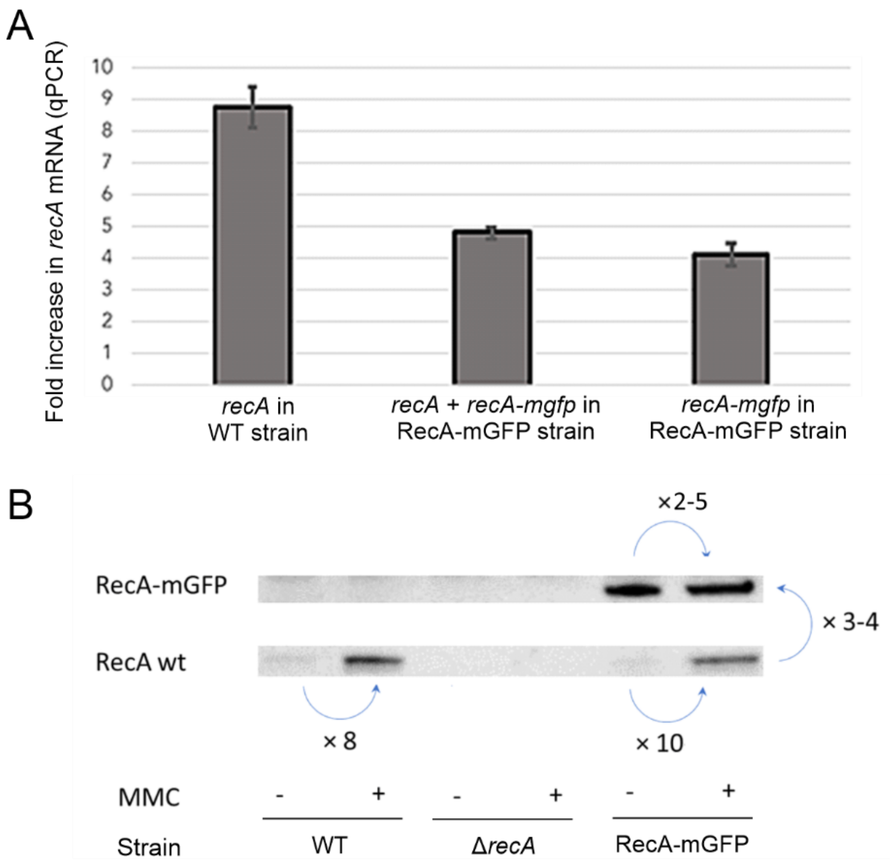


**Figure S4.** A) qPCR and B) Western blot results show both *recA* and *gfp* are upregulated on addition of MMC. The *recA* mRNA levels increase approximately 9 fold upon treatment with MMC in the wild type MG1655 strain (WT), as mirrored by 8-fold increase of RecA protein in the blot in the wild type and merodiploid strains. The merodiploid labeled strain shows slightly less induction upon MMC treatment – there is a 5 fold increase in total *recA* mRNA (*recA* + *recA-GFP*) and about 4 fold increase in *recA-GFP* only mRNA upon mitomycin C treatment, as mirrored by a 2-5-fold increase in RecA-GFP protein. Bars indicate the average of n=3 qPCR replicates with standard errors. The Western blots imply a ratio of at least an order of magnitude greater RecA-mGFP over wild type RecA levels before MMC treatment, and a ratio of around 3-4:1 following treatment. From the these ratios we infer approximate correction factors of 1.0 ± 0.1 and 1.3 ± 0.1 respectively to recover estimates of the total *in vivo* amounts of wild type RecA + RecA-mGFP protein from the measured *in vivo* amounts of RecA-mGFP protein.

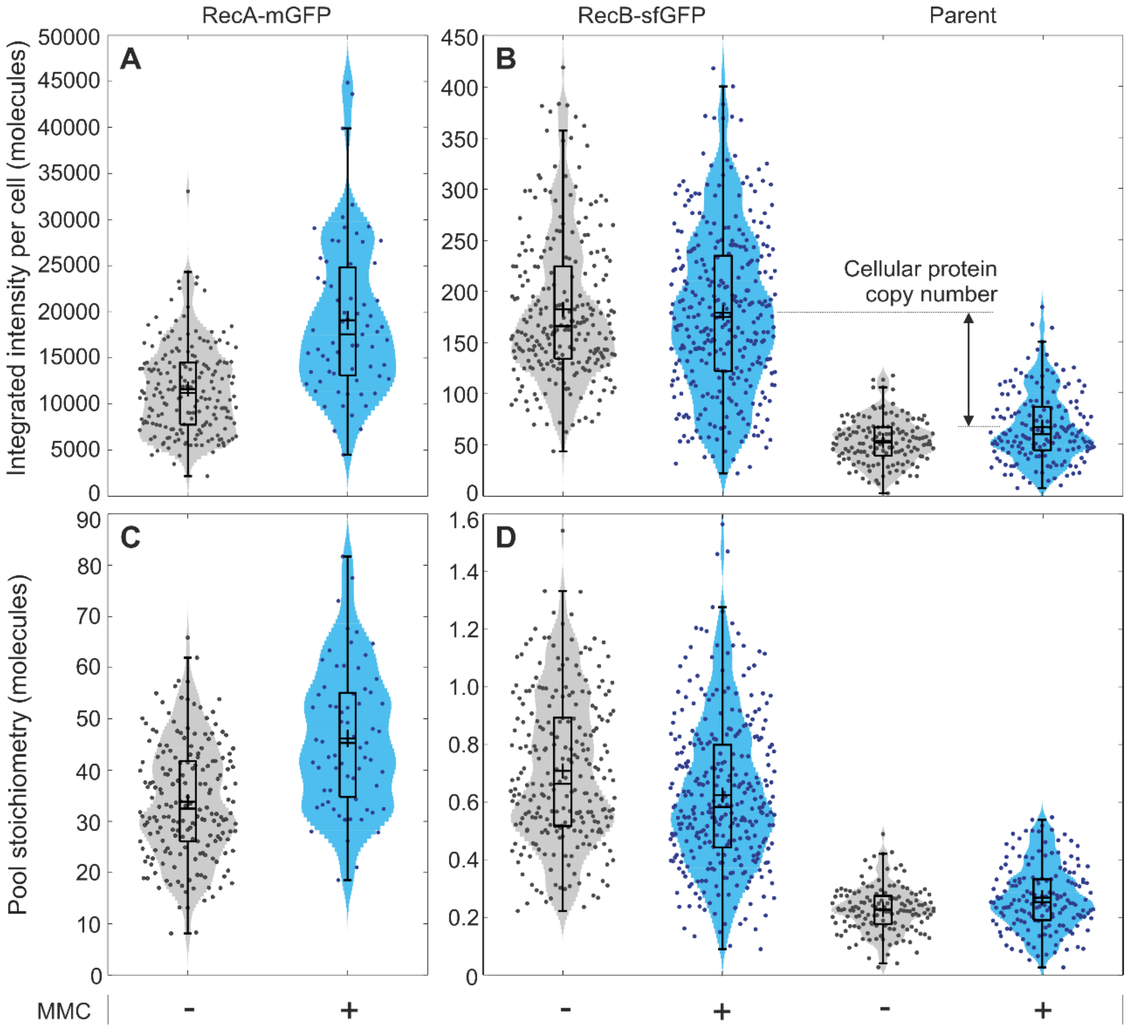


**Figure S5.** Population-level distributions of integrated intensities of each cell expressed in molecules (GFP equivalents) of A) RecA-mGFP protein and B) RecB protein *vs.* the unlabeled MG1655 parental wild type strain as determined from total fluorescent intensity. Individual cell data are shown overlaid with the associated 'violin' kernel density distributions and boxplots with interquartile range (box), median (horizontal solid line) and mean with standard error (black cross). Cellular protein copy numbers are estimated from the differences between mean values of integrated intensity of labeled and parental strains under the corresponding MMC condition. By determining the mean concentration of protein outside tracked foci, we find distributions of the upper possible limit of untracked stoichiometry of C) RecA-mGFP and D) RecB-sfGFP in the intracellular pool, vs. that of the unlabeled parent strain. Both copy and pool stoichiometry metrics are shown for cultures with (blue) and without (grey) MMC treatment. Statistics as described per condition in Table S1 in the range of 67 - 307 cells per strain.

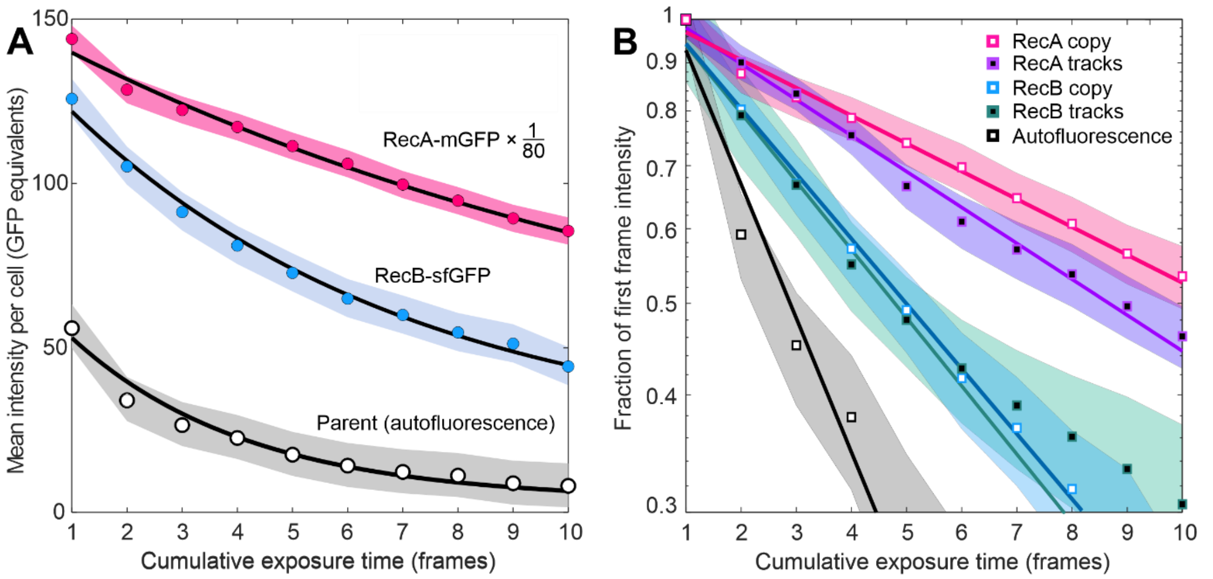


**Figure S6.** A) Photobleaching traces expressed as apparent integrated intensity of RecA-mGFP (multipled by a factor of 1/80 solely for clarity of graphical display to enable simple comparisons by eye with the other traces), RecB-sfGFP and the autofluorescent parent strain over initial exposure timepoints in the absence of MMC. Not only are these distinguishable in magnitude, but they photobleach at different initial rates. Shaded areas are 95% confidence intervals on data at each timepoint. Solid lines shown are single exponential fits to first 10 points unconstrained in height and slope, but with fixed baselines determined after 50 frames. B) Normalised photobleaching traces from cellular protein copy number, or from mean track brightness over time. The same fits are shown on a logarithmic scale to compare rates more readily. The decay of the excess RecB signal strongly resembles that of RecB confined to tracks; this indicates correctly labeled RecB-sfGFP is the dominant contribution to the apparent RecB copy rather than increased autofluorescence. Both A) and B) derive from the same MMC- datasets: either N=190 RecA-mGFP cells containing n=316 tracks, or N=249 RecB cells containing n=514 tracks, or N=103 parental MG1655 cells.

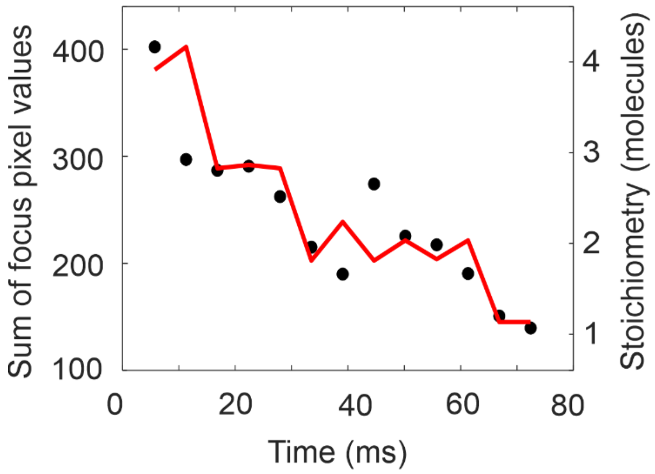


**Figure S7.** Stepwise photobleaching to determine characteristic number of counts per GFP per frame. A single track of RecA-mGFP is shown with the sum of pixel values in each of the constituent foci in subsequent frames over real time (circles) and the edge-preserving Chung-Kennedy filtered trace, window width 3 frames (line).

### Supplementary methods

*Segmentation and image post-processing*

An in-house manual segmentation protocol in MATLAB was used to identify individual cells from widefield images. This step was key to providing whole-cell statistics such as numbers of tracks per cell (Figure 1), and cellular protein copy number (Figure S5), and key to excluding background noise in all other measures. To improve throughput and reproducibility, we also developed a complementary AI-driven method to segment cell-containing image pixels from background pixels. This method consisted of a standard U-Net convolutional neural network with four layers [92], followed by object labelling using CellProfiler software [93]. The U-Net was trained with a binary cross entropy loss function [94] and an Adam optimizer [95] for 50 epochs (a machine learning term referring to the training iteration) with 10% dropout for the top two layers, 20% dropout for the bottom two layers, and a batch size of 6. We used 248 training images with a 9:1 training:validation split and reserved 76 images for testing. Images were prepared by registration of the brightfield and fluorescence micrographs and were constrained to be 256x256 pixels in size, roughly the size of our epifluorescence excitation spot. The best performance was obtained with a learning rate of $2.5 \times 10^{-4}$ and a patience value of 7 epochs to avoid overfitting [96], giving an accuracy of 0.995 for all image pixels on our validation and testing sets (Figure S2). For real-life tests we compared our U-Net segmentation of fields of view with varying densities of cells with MATLAB hand-segmentations of the same image which were used as ground truth (reliable known standard). We find that in cases of low cell number density, the segmentation classifies all pixels with good (>95%) accuracy. As cell number density increases, the pixel classification accuracy decreases (towards ~70%) due to ambiguity at cell boundaries and contacts. This trend is to be expected since the brightfield training sets include primarily well-separated *E. coli* cells at low contrast. Our choice of ground truth is not itself immune to this ambiguity at cell contacts, which at the cost of convenience, would be better specified with a membrane-localized fluorescent control. Nonetheless, this proof-of-concept implementation is more effective than pretrained or unsupervised methods for the Slimfield modality to date. To achieve a fully automated pipeline with true object labelling, we will incorporate additional postprocessing steps such as denoising, background subtraction, and removal of spurious cell boundaries [97].
